# Mechanisms of motor-independent membrane remodeling driven by dynamic microtubules

**DOI:** 10.1101/590869

**Authors:** Ruddi Rodríguez-García, Vladimir A. Volkov, Chiung-Yi Chen, Eugene A. Katrukha, Natacha Olieric, Amol Aher, Ilya Grigoriev, Magdalena Preciado López, Michel O. Steinmetz, Lukas C. Kapitein, Gijsje Koenderink, Marileen Dogterom, Anna Akhmanova

**Author notes:** Lead contact: Anna Akhmanova.

## Abstract

Microtubule-dependent organization of membranous organelles, such as the endoplasmic reticulum, occurs through motor-based pulling and by coupling microtubule dynamics to membrane remodeling. How highly transient protein-protein interactions occurring at growing microtubule tips can induce load-bearing processive motion is currently unclear. Here, we reconstituted membrane tubulation in a minimal system with giant unilamellar vesicles, dynamic microtubules, End-Binding (EB) proteins and a membrane-targeted protein that interacts with EBs and microtubules. We showed that these components are sufficient to drive membrane remodeling by three mechanisms: membrane tubulation by growing microtubule ends, motor-independent membrane sliding along microtubule shafts and pulling by shrinking microtubules. Experiments and modeling demonstrated that the first two mechanisms can be explained by adhesion-driven biased membrane spreading on microtubules. Force spectroscopy revealed that attachments to growing and shrinking microtubule ends can sustain forces of ∼0.5 and ∼5 pN, respectively. Rapidly exchanging molecules that connect membranes to dynamic microtubules can thus bear sufficient load to induce membrane deformation and motility.

## Introduction

Membrane-bound cellular organelles can acquire complex shapes, which are essential for their functions. Defects in organelle morphology and distribution can affect cell viability and lead to human diseases (Area-Gomez and Schon, 2016; Fu and Holzbaur, 2014; Westrate et al., 2015). Formation of membrane structures with a high curvature, such as tubules, is energetically unfavorable and thus requires applied forces (Helfrich, 1973; Helfrich and Servuss, 1984; Jarsch et al., 2016; McMahon and Boucrot, 2015; Zimmerberg and Kozlov, 2005). In cells, the generation and stabilization of curved membranes depends on membrane-deforming proteins and the cytoskeleton (Jarsch et al., 2016; McMahon and Boucrot, 2015; Simunovic et al., 2016; Zimmerberg and Kozlov, 2005).

Microtubules (MTs) are major cytoskeletal filaments, and the forces generated by MTs are important for many cellular processes (Brouhard and Rice, 2018; Dogterom et al., 2005). Dynamic MTs generate pushing and pulling forces with the magnitude of several piconewtons (Dogterom and Yurke, 1997; Grishchuk et al., 2005). Pushing forces are observed when a MT polymerizes against an obstacle such as a membrane; these forces can be used to position intracellular structures as well as whole MT networks like the mitotic spindle (Dogterom et al., 2005; Garzon-Coral et al., 2016). Pulling forces in turn are generated through protein attachment to depolymerizing MTs; these forces play a crucial role in chromosome separation during cell division but can also have other functions (Koshland et al., 1988; McIntosh et al., 2010; Waterman-Storer et al., 1995). Finally, MT-based molecular motors, kinesins and dyneins, can generate force to position and shape cellular organelles (Vale, 2003).

The endoplasmic reticulum (ER) is a complex membrane structure, which adopts a variety of morphologies ranging from the nuclear envelope to dynamic tubules. These morphologies depend on specific membrane-shaping proteins (Powers et al., 2017) and on the interactions with the cytoskeleton (Westrate et al., 2015). In particular, several types of MT-based processes have been implicated in the extension and positioning of ER tubules. ER tubules can extend along pre-existing MTs in a process termed sliding (Figure 1A), which can be driven by MT-based molecular motors (Friedman et al., 2010; Grigoriev et al., 2008; Waterman-Storer and Salmon, 1998; Woźniak et al., 2009). The extraction of membrane tubes by MT motors that move on stabilized MTs has been extensively studied by *in vitro* reconstitution experiments (Campàs et al., 2008; Koster et al., 2003; Leduc et al., 2004; Roux et al., 2002; Shaklee et al., 2008). However, it is still not clear how MT-based motors attach to the ER and how much they contribute to ER tubulation in cells.

**Figure 1.**
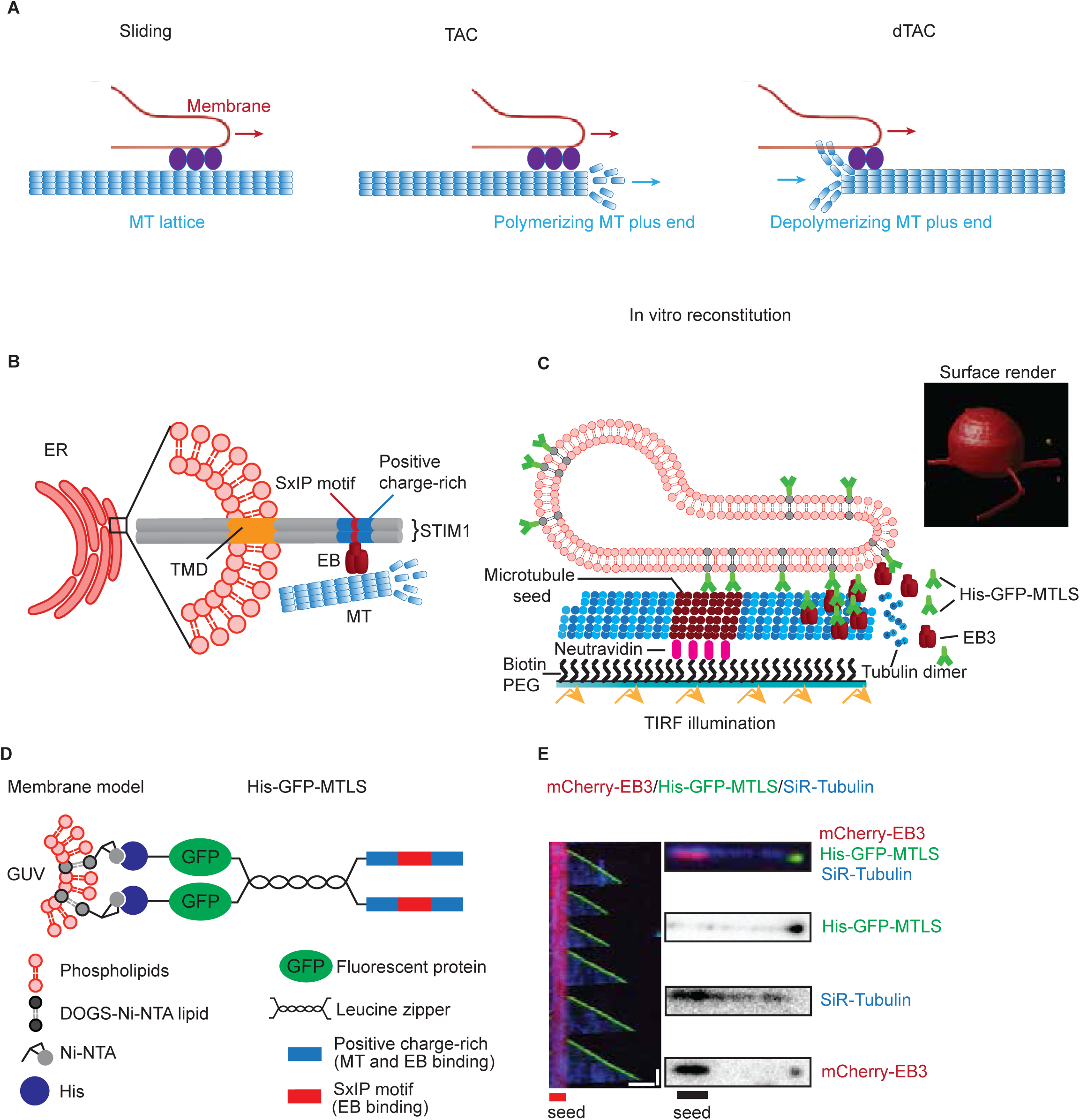
MT-driven formation of membrane tubes. (A) A scheme of the processes implicated in the extension and positioning of ER tubules. (Left) ER tubules can extend along pre-existing MTs in a process termed sliding, which can be driven by MT-based molecular motors. (Center) An ER tubule can be formed when a membrane attaches to the plus end of a growing MT via a membrane-MT tip attachment complex (TAC). (Right) ER tubules can be pulled by depolymerizing MT ends (dTAC) (B) A scheme of a TAC formed on the ER membranes by STIM1 and an EB protein. (C) A scheme of the assay that includes GUVs, GMPCPP-stabilized MT seeds (which contain some biotinylated tubulin and are attached to biotin-PEG-coated glass slides through neutravidin), unlabeled tubulin, mCherry-EB3 and His-GFP-MTLS that binds to DOGS-NTA-Ni on the GUV surface. (Right) Surface render of a confocal Z stack obtained from a GUV after the incubation with dynamic MTs in presence of 200 nM mCherry-EB3 and 15 nM His-GFP-MTLS. Confocal Z stacks were obtained using spinning disk microscope. Scale bar: 5 μm. (D) Schematic representation of His-GFP-MTLS, used as a membrane-MT linker in this study. MTLS (MT Tip Localization Sequence) consists of a 4-amino acid motif SxIP embedded in an intrinsically disordered, positively charged polypeptide region. (E) Three color TIRFM kymographs and snapshots, showing the accumulation of mCherry-EB3 (red) and His-GFP-MTLS (green) at the plus end of a growing MT (cyan) grown from rhodamine-tubulin labeled MT seeds. Kymograph scale bars: horizontal, 2 μm, vertical, 1 min. Snapshots, scale bar: 1 μm. See also Figure S1A-E, and Video S1.

An ER tubule can also be formed when a membrane attaches to the plus end of a growing MT via a membrane-MT tip attachment complex (TAC) (Grigoriev et al., 2008; Guo et al., 2018; Waterman-Storer et al., 1995; Waterman-Storer and Salmon, 1998) (Figure 1A). Experiments in cells have demonstrated that TAC formation requires two types of MT plus-end tracking proteins (+TIPs): End Binding (EB) proteins, which recognize the stabilizing cap at growing MT ends (Maurer et al., 2012), and the ER-resident transmembrane protein STIM1 (Grigoriev et al., 2008). STIM1 specifically accumulates at the plus ends of growing MTs in an EB-dependent manner, because it contains an EB-binding MT Tip Localization Sequence (MTLS), which consists of four amino-acid long motif SxIP embedded in an intrinsically disordered, positively charged polypeptide region (Figure 1B,D) (Honnappa et al., 2009). Due to the presence of positively charged sequences, STIM1 and other proteins containing an MTLS can interact with MTs not only though EBs, but also directly, by binding to the negatively charged MT surface (Honnappa et al., 2009). However, it is currently unknown whether the two known TAC components, an EB protein and a membrane associated EB partner, are sufficient for membrane deformation by dynamic MTs without the participation of molecular motors. It is also unclear whether and how various +TIPs, most of which arrive to growing MT tips by diffusion and rapidly exchange at these ends (Bieling et al., 2007; Dragestein et al., 2008), can mediate forces that induce membrane tube formation and promote processive tube extension.

Finally, recent work has shown that ER tubules can be pulled by depolymerizing MT ends, a mechanism termed dTAC (Guo et al., 2018)(Figure 1A). The molecular mechanisms underlying dTAC are currently unknown and it is unclear whether dTACs can be driven by the same molecules that are responsible for TAC formation.

Here, we show that membrane tubulation by sliding, TAC and dTAC mechanisms can be reconstituted in a minimal *in vitro* system containing dynamic MTs, an EB protein and an engineered membrane-bound protein that interacts with MTs and EBs. Experiments with optical tweezers demonstrated that the magnitude of the force generated by TAC- and dTAC-mediated mechanisms is in the range of 0.5 and 5 pN, respectively. Motor-independent membrane sliding and TAC-based membrane remodeling can be explained by adhesion-driven spreading of membranes on MTs. The combination of experiments and modeling described here thus reveals that the forces generated by MT dynamics in combination with rapidly exchanging MT-associated proteins, with no participation of molecular motors, can induce robust membrane tubulation and can make a major contribution to shaping endomembranes.

## Results

### A membrane-bound MTLS-containing protein and EB3 promote membrane tube extension by dynamic MTs

To investigate the mechanism of force generation underlying TAC-based membrane extension, we reconstituted the interaction between membranes and dynamic MTs in a system with a minimal set of components (Figure 1C). To this end, we used giant unilamellar vesicles (GUVs) prepared from POPC (94.95%), DOGS-NTA-Ni (5%) and Rh-PE (0.05%). The GUVs were produced by swelling a dried film of lipids in a 300 mM sucrose solution. To avoid membrane tube formation due to the osmotic stress, we adjusted the osmolarity of the solution outside of the GUVs to 320 mM, so that the difference in osmolarity inside and outside of the GUVs would not exceed 10%. MTs were grown from GMPCPP-stabilized MT seeds as described previously (Aher et al., 2018; Bieling et al., 2007) in the presence of tubulin, mCherry-EB3 and an engineered protein that was termed His-GFP-MTLS (Figure 1D), which could interact with the GUVs, EB3 and MTs. HIS-GFP-MTLS contains the C-terminal 43 residues of human MT-actin cross-linking factor 2 (MACF2) bearing an EB-binding SxIP motif embedded in an intrinsically disordered, positively charged region (Honnappa et al., 2009). Due to the presence of positive charges, MTLS has some affinity for the surface of MTs even in the absence of EB proteins (see below). His-GFP-MTLS was dimerized via the leucine zipper from GCN4 and tagged with the green fluorescent protein (GFP) (Figure 1D). As described previously (Honnappa et al., 2009), this well-characterized fusion protein is representative of other EB-dependent MT plus end ligands, such as STIM1, which is also a dimer with a single SxIP motif embedded in a positive charge-rich sequence exposed in the cytoplasm (Honnappa et al., 2009) (Figure 1B). A 6-Histidine tag (His) was added to the N-terminus of GFP for the interaction with the Ni-NTA moiety on the GUV surface.

MTs and GUVs were visualized using Total Internal Reflection Fluorescence (TIRF) or confocal microscopy (Figure 1C). In the assay with dynamic MTs, His-GFP-MTLS was strongly enriched at growing MT ends in an EB3-dependent manner, consistent with previous reports (Honnappa et al., 2009), and its accumulation at MT tips increased at higher EB3 concentrations. Furthermore, His-GFP-MTLS was weakly bound along MT shafts (Figures 1E, S1A-E, and Video S1), likely because MTs are negatively charged and the protein is positively charged; this binding was EB3-independent. In absence of GUVs, the amount of His-GFP-MTLS at the MT tip remained constant over time (Figure S1D). When GUVs were included in the assay, the number of His-GFP-MTLS molecules attached to the MT tip decreased with time, suggesting that most of the His-GFP-MTLS molecules were recruited to the GUV surface (Figure S1D). His-GFP-MTLS can thus potentially mediate the interaction between GUVs and MTs.

When GUVs were combined with MTs, within a few minutes, membrane tubes started to form (Figure 1C, surface render). These tubes co-localized with MTs, which were visualized using SiR-tubulin, a fluorescent probe based on the silicon-rhodamine and the MT-binding drug Docetaxel (Figure 2A), indicating that MTs supported membrane tube extension. Earlier studies reported generation of tubular networks when GUVs were incubated with stable MTs and motors (Koster et al., 2003; Leduc et al., 2004; Roux et al., 2002; Shaklee et al., 2008; Yamada et al., 2014). In contrast, our assay did not include motor proteins, demonstrating that a motor-independent MT-based mechanism can extract tubes from a lipid vesicle.

**Figure 2.**
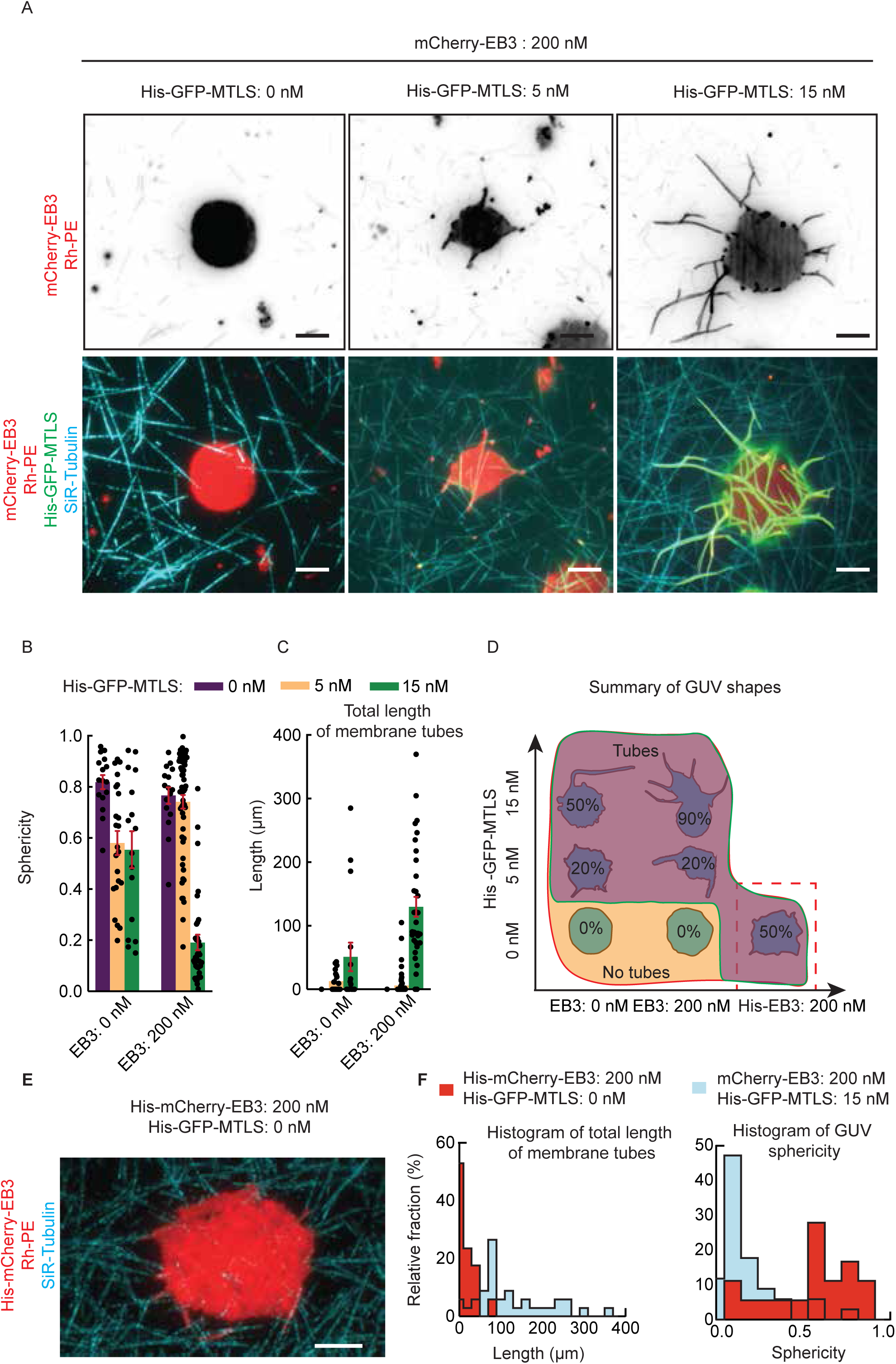
Formation of long membrane tubes in the presence of dynamic MTs, EB3 and His-GFP-MTLS. (A) Representative images of a tubular network (red) observed at different concentrations of His-GFP-MTLS and 200 nM EB3. Each image is the maximum intensity projection obtained from 100 frames. Scale bar: 5 μm. (B, C) GUV sphericity (B) and the total length of membrane tubes (C), measured at different His-GFP-MTLS and EB3 concentrations. Without EB3, 0 nM MTLS, n=16 GUVs; 5 nM MTLS, n=23 GUVs; 15 nM MTLS, n=16 GUVs. At 200 nM EB3, 0 nM MTLS, n=16 GUVs; 5 nM MTLS, n=57 GUVs; 15 nM MTLS, n=34 GUVs. Error bars indicate SEM. See also Figure S2. (D) A diagram showing characteristic GUV shapes observed as a function of His-GFP-MTLS and EB3 concentrations. The numbers represent the proportion of tubular GUVs observed without EB3 at 0 nM MTLS, n=16 GUVs; 5 nM MTLS, n=23 GUVs; 15 nM MTLS, n=16 GUVs. At 200 nM EB3, 0 nM MTLS, n=16 GUVs; 5 nM MTLS, n=57 GUVs; 15 nM MTLS, n=34 GUVs. (E) Still image of a GUV (red) in the presence of dynamic MTs (blue) and 200 nM His-EB3. (F) Histogram of total length of membrane tubes (left) and GUV sphericity (right) measured at 200 nM of His-EB3 without His-GFP-MTLS, n=34 GUVs (red) and at 200 nM of mCherry-EB3, 15 nM of His-GFP-MTLS, n=18 GUVs (blue).

We next investigated the effect of EB3 and His-GFP-MTLS on membrane tube formation. To characterize membrane morphology, we measured the GUV sphericity and the total length of all tubes in the network formed by each vesicle (Koster et al., 2003; Legland et al., 2016)(Figures 2A-C and S2). In the absence of His-GFP-MTLS, GUVs preserved their round shape even at a high (200 nM) EB3 concentration, and the transition to the tubular regime did not occur (Figure 2A-D). In contrast, in the presence of His-GFP-MTLS and in the absence of EB3, some tubes were observed and the number of detected tubes increased with the His-GFP-MTLS concentration (Figure 2A,C,D). Short tubes could be detected at 200 nM EB3 and 5 nM His-GFP-MTLS (Figure 2A,C,D); however, only when the concentration of His-GFP-MTLS was increased to 15 nM in the presence of 200 nM EB3, extensive tubulation of 90% of the GUVs was observed (Figure 2A-D). These data indicate that MT-driven membrane tubulation can be reconstituted using proteins that couple membranes to dynamic MTs.

To test whether MT-membrane interaction can be supported by EB3 alone, we used its 6-Histidine-tagged version (His-EB3) that could directly interact with growing MT tips and DOGS-NTA-Ni on the GUV surface. However, even at 200 nM His-EB3, only short tubular structures were present on 50% of the GUVs (Figure 2D-F), likely because the affinity of EB3 for MT shafts is low. These results indicate that an EB3 protein directly linked to membranes can promote some membrane tubulation, but the generation of long membrane tubes occurs much more efficiently when an EB3 protein is combined with a MT-binding membrane-attached protein.

### MT-dependent membrane tubulation is limited by tension

Previous studies have shown that ligand-dependent adhesion of GUVs to the surface of a solid substrate or to a nanofiber triggered the formation of membrane tubules (Charles-Orszag et al., 2018; Feder et al., 1995; Lobovkina et al., 2010). When a GUV faces an adhesive substrate in the presence of mobile ligands, the ligands that bind to the substrate move to the contact region and increase the adhesion strength (Brochard-Wyart and de Gennes, 2002; Bruinsma and Sackmann, 2001). We investigated whether His-GFP-MTLS was enriched in the regions where GUVs interacted with MTs and found that the average fluorescence intensity of the His-GFP-MTLS on MTs bound to the membrane was indeed higher than on free MTs or membranes that were not in contact with MTs (Figure 3A,B). His-GFP-MTLS proteins thus converged towards the sites of MT-membrane contact, suggesting an increase in the number of bonds between the two structures.

**Figure 3.**
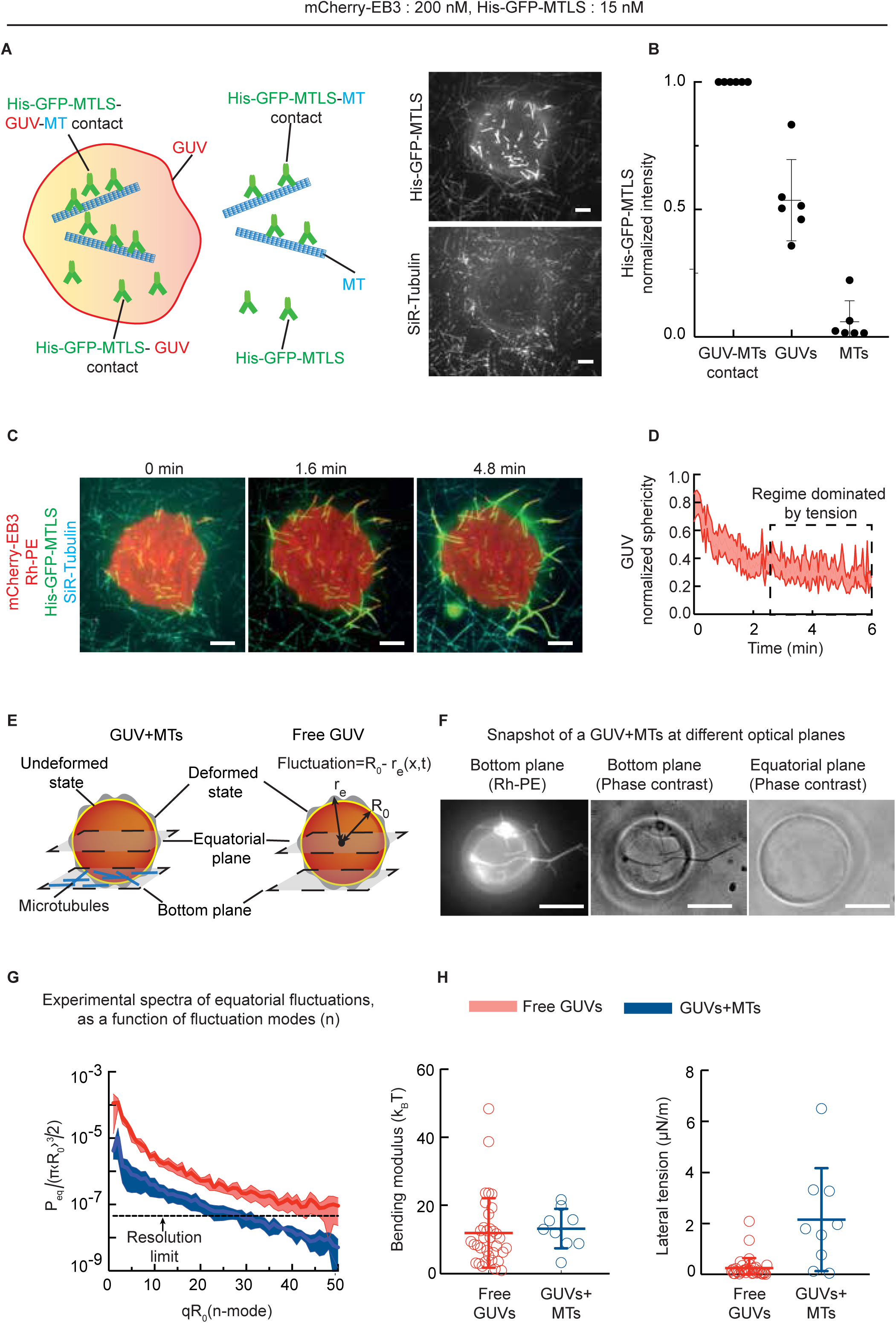
Control of MT-dependent formation of tubular membrane networks by adhesion and tension. (A) Left: Schematic representation of different binding states of His-GFP-MTLS in the assay. Right: Snapshot of a TIRFM time lapse movie showing free MTs and MTs bound to a GUV. Only the His-GFP-MTLS and MT channels are shown. Scale bar: 5 μm. (B) Averaged MT intensity profiles of the His-GFP-MTLS channel for MTs attached to a GUV (n = 6 movies 62 MTs), the region of the GUV close to the attached MT (n = 6, movies, 62 profiles) and free MTs (n=6 movies, 106 MTs). The values were normalized to the mean values of the MTs attached to a GUV. (C) Snapshots of a TIRFM time lapse movie showing the development of a tubular membrane network over time. Scale bar: 5 μm. See also Video S2 (D) Averaged values of GUV sphericity as a function of time, normalized to the maximum experimental value. The shaded area represents SEM (n=11). (E) Schematic representation of the two conditions where membrane fluctuations were investigated, GUVs in the presence of MTs and free GUVs. The equatorial and the bottom plane are indicated; blue lines represent MTs. The membrane fluctuates around an equilibrium position (undeformed state, yellow line) defined by the time-average of the GUV radius R_0_. Fluctuations at each point (deformed state) are represented in grey and are defined by local deviations of the equatorial radius r_e_ from the average radius R_0_. (F) Phase contrast and fluorescence images of a GUV interacting with MTs. Bottom plane is shown on the left and center, equatorial plane is on the right. Scale bar: 5 μm. (G) Experimental fluctuation spectrum calculated from the time-averages of quadratic fluctuation amplitudes of the equatorial modes, for free vesicles (red, n=39) and vesicles in contact with MTs (blue, n=13), as a function of the fluctuation mode *n*. The averaged curves were calculated after correcting each individual spectrum for pixelization noise and divided by π〈*R_0_*〉*^3^/2*. The black dashed line represents the resolution limit, estimated by measuring the fluctuation spectrum of a fixed object in the focal plane. Shaded areas represent SEM. See also Figure S3. (H) Estimated values of the bending modulus and lateral tension for free GUVs (n = 38) and GUVs in contact with MTs (n = 10). Error bars indicate SD.

Next, we investigated the dynamics of formation of tubular membrane networks by measuring GUV sphericity (Legland et al., 2016) over time. At the beginning of the assay, sphericity quickly decreased, indicating that the network of membrane tubes rapidly expanded (Figure 3C,D). At later time points, the sphericity value became almost constant as tube extension slowed down (Figure 3C,D and Video S2). We hypothesized that this was caused by the increase in lateral tension after the excess of membrane area was used up due to membrane redistribution into tubes (Cuvelier et al., 2005; Evans and Rawicz, 1990; Fournier et al., 2001; Sengupta and Limozin, 2010). To test this possibility, we characterized thermal membrane fluctuations, which are controlled by the balance between thermal energy, lateral tension and membrane stiffness (Helfrich, 1973; Helfrich and Servuss, 1984). Flickering spectroscopy (Pecreaux et al., 2004; Rodriguez-Garcia et al., 2009) was used to analyze membrane undulations in freely fluctuating GUVs and GUVs attached to MTs (Figure 3E). Fluctuations of the vesicle radius were recorded at the equatorial plane by video microscopy (Figure 3E, F) and expanded in series of discrete Fourier modes *n*=2,3…50. We then calculated the time-average of quadratic fluctuation amplitudes. The experimentally estimated amplitudes were corrected for the noise caused by the finite spatial resolution of the imaging (pixelization) and plotted as a function of the mode *n*, known as a fluctuation spectrum (FS)(Figures 3G and S3). In the presence of MTs, the amplitude of membrane fluctuations was reduced by an order of magnitude (Figure 3G), with amplitudes lower than our detection limit at high fluctuation modes. The elastic modulus that we measured (Figure 3H) was in good agreement with previously published values (Rodriguez-Garcia et al., 2009), indicating the similarity in membrane stiffness in both conditions. Importantly, lateral membrane tension was significantly higher in the presence of MTs, leading to the reduction in the amplitude of fluctuations (Figure 3G, H). Taken together, our data indicate that GUV extension along MTs follows previously described membrane adhesion dynamics (Bernard et al., 2000; Boulbitch et al., 2001; Cuvelier and Nassoy, 2004; Sengupta and Limozin, 2010), where the initial spreading is counteracted by lateral membrane tension.

### Membrane remodeling occurs by three mechanisms

To understand better how membrane tubes are pulled in our assay, we investigated their dynamics in more detail. We observed some membrane tubes sliding along MT shafts (Figures 4A, S4A and Video S3). Tube sliding occurred with the speed of ∼5-8 µm/min, faster than the elongation rate of a growing MT end, which did not exceed ∼4 µm/min in most of our assays (Figures 4B-D and S4B). When the tip of a sliding membrane tube reached the end of a growing MT, the membrane slowed down because it could not extend beyond the MT tip and its spreading was thus limited by the speed of MT growth (Figure 4B,C). A growing MT end could also initiate tube formation without prior sliding along a pre-existing MT (Figure S4C and Video S4). As before, in presence of EB3, we observed an enrichment of the His-GFP-MTLS at the MT tips (Figures 4C, S4D and Video S4). Interestingly, the interaction of a membrane with a MT end correlated with a decreased rate of MT growth (Figure 4D). One potential reason for this effect is the decrease in tubulin diffusion to the MT tip caused by membrane shielding. This conclusion is supported by the observation that a similar effect was observed also in the absence of His-GFP-MTLS (Figure 4D), in conditions where there was no direct interaction between the MT and the membrane.

**Figure 4.**
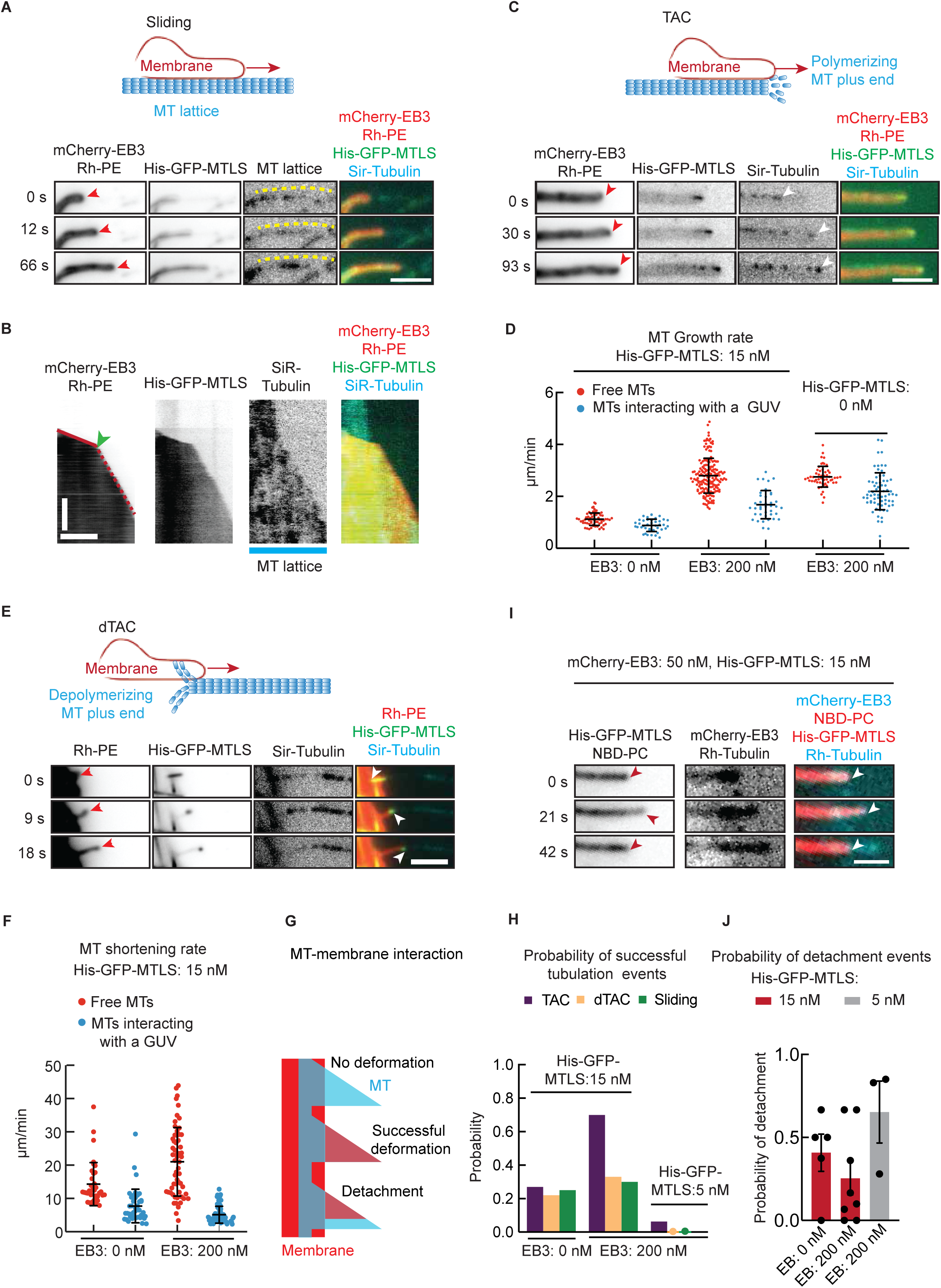
Three mechanisms of MT-induced membrane tube formation. (A,C) Time lapse images of a membrane tube (red arrowheads) sliding along a MT (dashed yellow line) (A), or moving together with the plus end of a growing MT (C)(white arrowheads) in the presence of 200 nM mCherry-EB3 (A), 50 nM mCherry-EB3 (C) and 15 nM His-GFP-MTLS. Scale bar: 1 μm. See also Figure S4A,S4C,S4D,S4F and S4G. Video S3 and Video S4. (B) Left: Kymograph of a membrane tube sliding along a MT and catching up with the MT tip (green arrowhead). Scale bars: horizontal, 2 min, vertical, 2 μm. Right: Speed of membrane sliding at 0 nM (n=9), and 200 nM (n=12) EB3 and 15 nM His-GFP-MTLS. Error bars indicate SD. See also Figure S4B. (D) The growth rate of free MTs (red) and MTs in contact with a membrane (blue), at the indicated EB3 and His-GFP-MTLS concentrations; n, from left to right: 63, 41, 179, 40, 41 and 67. Error bars indicate SD. (E) Time lapse images of a membrane tube (red arrowhead) attached to the plus end of a depolymerizing MT (white arrowhead) (dTAC) in the presence of 15 nM His-GFP-MTLS without mCherry-EB3. Scale bar: 2 μm. See also Figure S4E and Video S5 (F) The shortening rate of free MTs (red) and MTs pulling a membrane (blue), at the indicated EB3 and His-GFP-MTLS concentrations; n, from left to right: 36, 41, 58 and 73. Error bars indicate the SD. (G) Schematic representation of different membrane deformations induced by dynamic MTs. (H) Probability of successful membrane tube generation events occurring through different mechanisms at 15 nM of His-GFP-MTLS. The probability of successful tubulation events was determined by dividing the number of observed tubes by the total number n of MT-membrane contacts that could potentially support a specific event (e.g. total number of MT-membrane contacts for sliding, membrane contacts with growing MTs for TACs and shrinking MTs for dTAC). 0 nM EB3:TAC (n=48 events, 7 experiments), dTAC (n=36 events, 5 experiments), sliding (n=50 events, 5 experiments). 200 nM EB3: TAC (n=68 events, 9 experiments), dTAC (n=6 events, 5 experiments), sliding (n=30 events, 5 experiments). 200 nM EB3 and 5 nM of His-GFP-MTLS, TAC (n=37 events, 3 experiments), dTAC (n=20 events, 3 experiments), sliding (n=41 events, 3 experiments) (I) Snapshots of a TIRFM movie showing a membrane tube (red arrowheads) detaching from a MT tip (white arrowheads). Scale bar: 1 μm. (J) Probability of detachment events from a MT tip. 0 nM EB3, n=50 events, 7 experiments; 200 nM EB3 n=68 events, 9 experiments, both in presence of 15 nM of His-GFP-MTLS; 200 nM EB3 and 5 nM of His-GFP-MTLS, n=37 events, 3 experiments. Error bar indicates SEM.

Recent work using structured illumination microscopy in living cells demonstrated that ER tubules could be pulled by depolymerizing MT ends (dTAC) (Guo et al., 2018). Such events were also observed in our assays, where shrinking MTs could extract long membrane tubes from GUVs (Figure 4E, S4E and Video S5). At the tips of dTAC-driven membrane tubes, we observed an accumulation of His-GFP-MTLS, but not of EB3 (Figures 4E, S4E and Video S5). Strikingly, the shortening rate of MTs that were in contact with a membrane was reduced ∼ 3 fold compared to the depolymerization rate of free MTs (Figure 4F). Shortening MT ends are characterized by rapid disassembly of tubulin protofilaments, which bend outwards and generate a power stroke. Protein complexes capable of following the ends of shortening MTs were shown to slow down the speed of MT disassembly, indicating a stable interaction with the disassembling MT end (Grishchuk et al., 2008; Volkov et al., 2015; Volkov et al., 2018). It is thought that proteins interacting with bent tubulin protofilaments can transmit force from a depolymerizing MT end to a cargo, causing its displacement (Koshland et al., 1988; Mitchison, 1988). Our results suggest that membrane-bound His-GFP-MTLS proteins accumulating at the interface between the depolymerizing MT and the membrane create an attachment site that can transmit the force generated by a shortening MT to the membrane.

Next, we estimated the probability of successful membrane tubulation events (Figure 4G) occurring through sliding, TAC and dTAC mechanisms at the contacts between the membrane and either MT shafts, growing MT ends or shrinking MT ends, respectively. In the absence of EB3, the probability of successful events was similar for all three mechanisms (Figure 4H). This could be expected, because all events depended exclusively on the membrane-MT linkages mediated by His-GFP-MTLS that were not sensitive to the state of the end or the lattice of a MT. However, whereas the probability of successful sliding and dTAC events was not affected by EB3, the probability of successful TAC events was much higher in the presence of EB3 (Figure 4H). This could be explained by EB3 preferentially concentrating at the growing MT ends, which in turn leads to a local increase in the MTLS accumulation and thus an increased efficiency of an extended force-generating contact (Figures 1D, 4C, S1A-C and S4D).

The interactions of GUVs with growing MT ends led to short (non-tubular) or long (tubular) membrane deformations (Figures 4C,S4F,S4G, and Video S4 and S6). These deformations retracted either because a MT depolymerized or because the membrane detached from the growing MT tip, similar to previous observations in cells (Grigoriev et al., 2008; Waterman-Storer et al., 1995; Waterman-Storer and Salmon, 1998) (Figures 4I, S4F,S4G, and Video S4 and S6). The fraction of MT-membrane contacts leading to the formation of tubes was higher if both His-GFP-MTLS and EB3 were present in the assay, and was increased at a higher His-GFP-MTLS concentration (Figure 4H). A higher concentration of His-GFP-MTLS also suppressed membrane tube detachment from MTs (Figure 4J) and thus promoted the formation of tubes. These data indicate that the increased abundance of the complexes formed by EB3 and His-GFP-MTLS at MT tips stimulates TAC-mediated membrane tubulation.

### Modeling of membrane spreading driven by TAC formation

We next set out to obtain a quantitative description of the bilayer spreading process by combining theory with experimental measurements of different parameters of membrane-MT interactions (Table 1). We first described the membrane sliding mechanism using a simple analytical model (Phillips et al., 2012; Van Kampen, 1992) (see Methods for details). We assumed that the interaction between GUVs and MTs is dominated by the binding of the His-GFP-MTLS to MTs. We modeled the formation of a fixed-size membrane-MT adhesion domain, where we assumed a constant number of His-GFP-MTLS-MT connections at the tip of a membrane extension (Figure 5A). The interactions between His-GFP-MTLS and MTs were described by a binding rate *k_on-m_* (Table 1, parameter 4) and a detachment rate *k_off-m_* (Table 1, parameter 3), which were calculated from the experimentally measured mean maximum spreading speeds of sliding membrane tubes and the analysis of single molecule binding events of the His-GFP-MTLS to MTs (Table 1, Figures 5A and S5A-F). The detachment rate from a MT was expected to increase with the force *k_off-m_ (F_m_)* (Bell, 1978; Erdmann and Schwarz, 2004; Evans and Calderwood, 2007), where *F_m_*, the force needed to pull a tube from a GUV depended of the elastic modulus and the lateral tension of the GUV (Derényi et al., 2002) (see Equation 7 in Methods). These values were measured by flickering spectroscopy (Table 1, parameters 8,9 and Figure 3G,H). In addition, we assumed that the membrane tension increased with tube length (see Equation 8 in Methods), and consequently, higher force was needed to keep a membrane tube extended when the tube length increased. The tip of the membrane deformation extended by a length of one tubulin dimer *d*=8 nm upon the formation of a new membrane-MT bond and decreased by the same distance if a bond at the membrane front was detached (Figure 5A and Methods).

**Figure 5.**
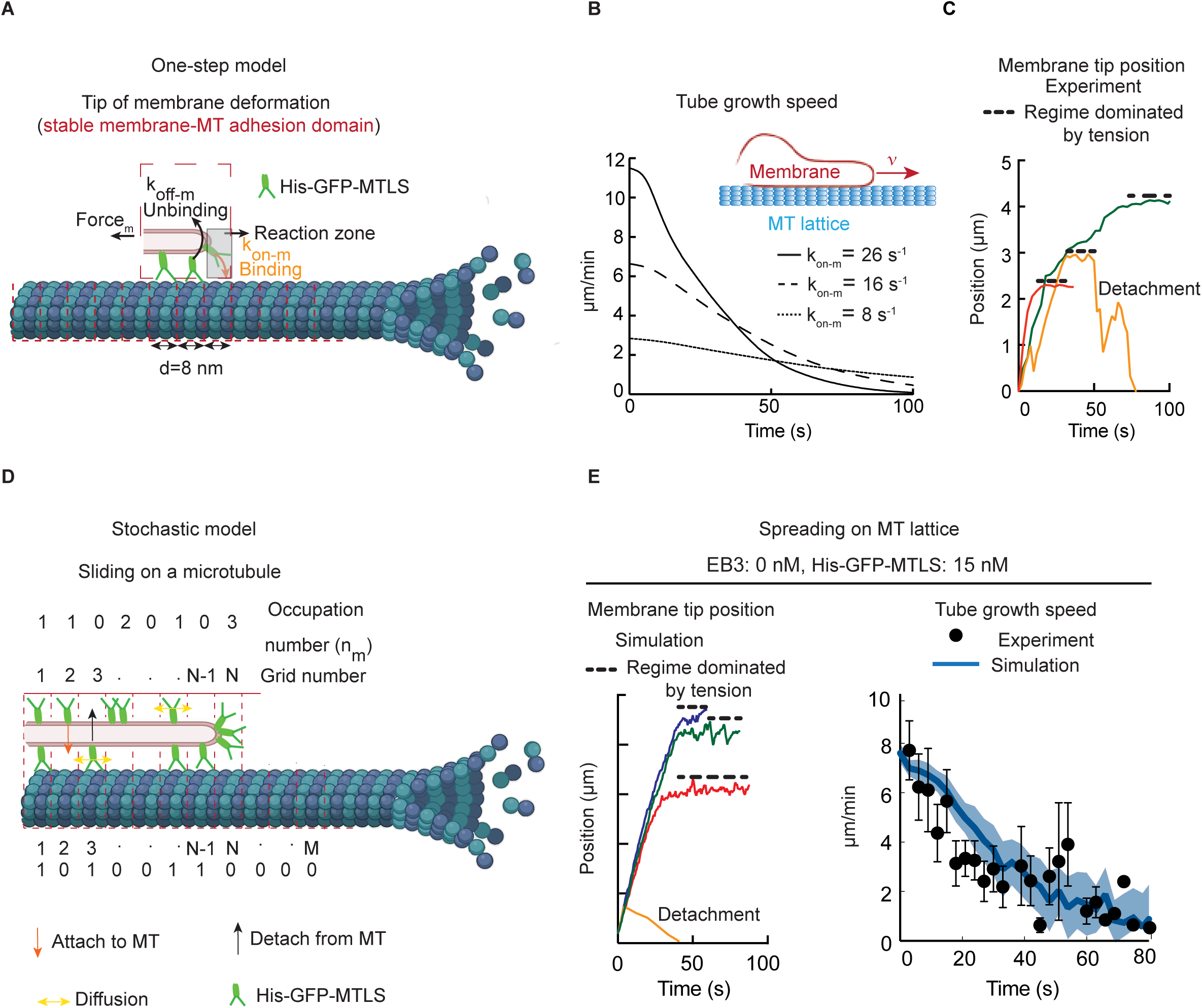
Modeling and simulations of membrane spreading along MT shafts and comparison of *in silico* and experimental data. (A) Kinetic scheme for the one-step model. (B) Predicted speed of membrane tip spreading along MT shafts as a function of time for three different values of the association rate using Equation 11 (Methods). Curves are averages of 100 individual traces with randomly sampled membrane parameters. (C) Experimental tip positions of membrane tubes sliding along MTs. Different traces are indicated with different colors. (D) Kinetic schemes for the simulations of a membrane sliding along a MT shaft. (E) Left: Simulated tip positions of membrane tubes sliding along MTs. Different traces of independent runs of the simulation are indicated with different colors. Right: Average tube velocity as a function of time (blue), estimated from 60 stochastic simulations of membrane spreading on MTs. The shaded region represents SEM. Experimental values of the average tube velocity are shown with black dots; error bars represent SEM, n=12.See also Figure S5A,B.

**Table 1:**
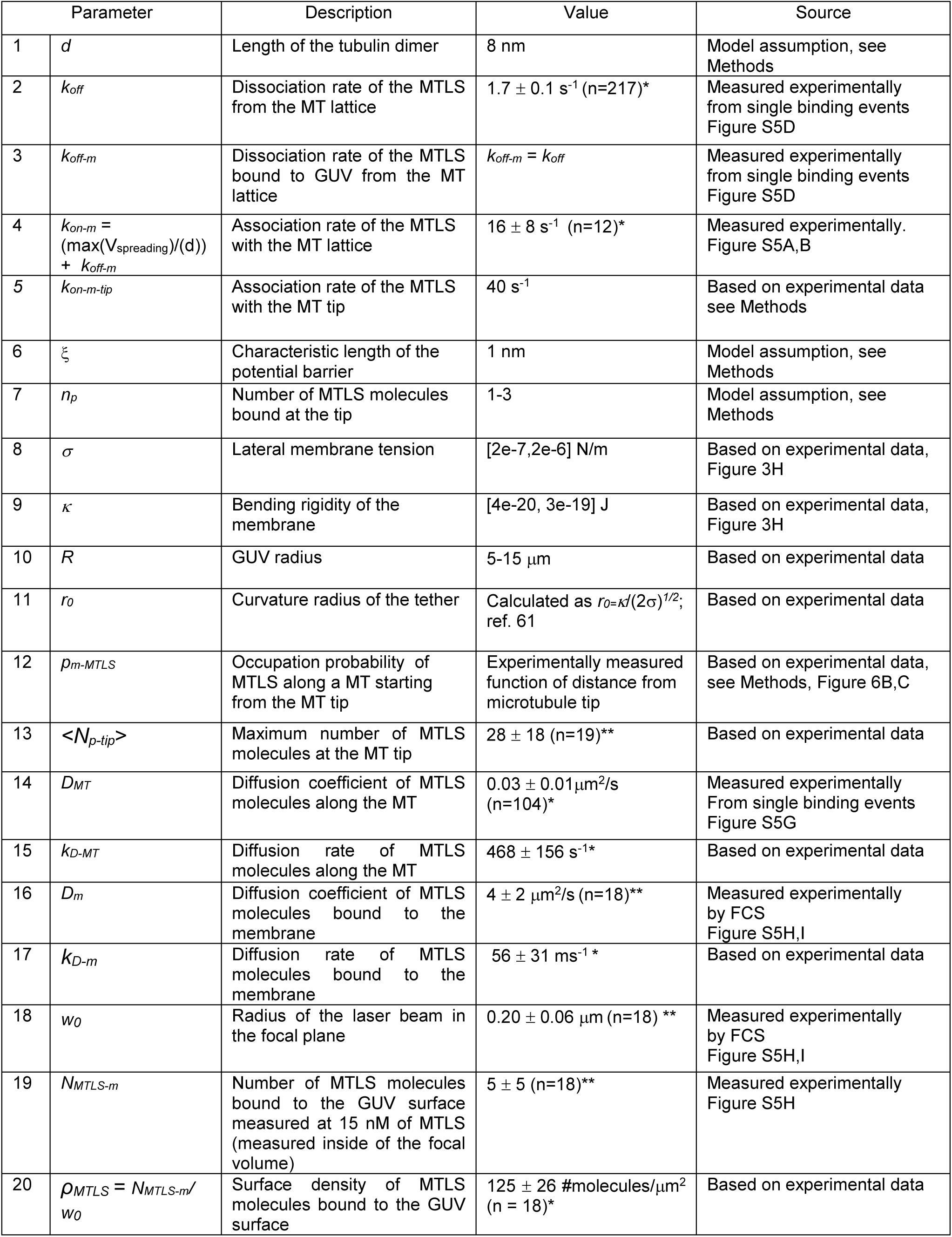
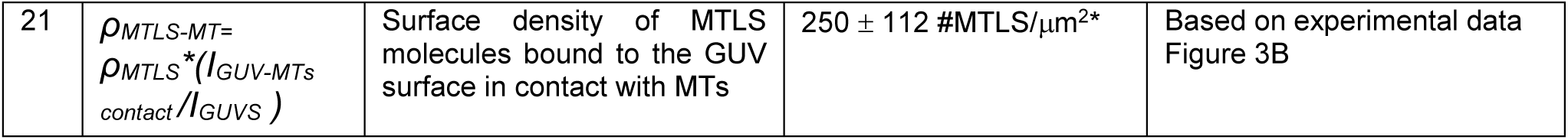
Parameters of the model and simulations. Dimeric HIS-GFP-MTLS protein (abbreviated as MTLS in the Table) was used for all measurements. *mean value ±SE,**mean value ± SD

In this model, the highest initial speed *V* of membrane spreading depends on the association rate *k_on-m_* and detachment rates *k_off-m_(F_m_)* as V*=d*[*k_on-m_-k_off-m_*(*F_m_*)] (Figure 5A, see Equation 11 in Methods). As the membrane tube increased in length, the force needed to pull a tube increases, and therefore the tube elongation rate slows down and finally drops to a value close to zero (Figure 5B). This final state matches well the saturation regime observed experimentally (Figure 5C).

To have a more realistic description of the tube elongation process that could be compared to experimental data, we next extended our simple model by taking into consideration the formation of bonds not only at the membrane tube tip but also all along the contact interface between the membrane deformation and a MT (Figure 5D, see Methods for more details)(Campàs et al., 2008). The position of the adhesion molecules changed with time due to the diffusion of the His-GFP-MTLS on the membrane and along the MT lattice (Figures 5D and S5G-I). In addition, we included in the model changes in MT length to reflect MT dynamics. Due to the fact that in this extended model a high number of reactions had to be considered, we performed *in silico* experiments using a stochastic approach (Gillespie, 1977, 2007) (see Methods for more details). When EB3 concentration is zero (Figure 5D), the estimated on-rate of MTLS is k_on-m_ = 16 s^-1^ (Table 1, parameter 4). In these conditions, simulations resembled the experimental observation quite well (Figure 5C, E left). The predicted sliding speed overlapped quite well with the experimental data obtained by measuring the spreading rate of membrane tubes on MT shafts (Figures 5E, right). In addition, the estimated membrane spreading speeds from simulations and experiments agreed with the predictions of the analytical model, showing a decay that was dominated by tension (Figure 5B, E, right).

Finally, we incorporated into the model the EB3-induced enrichment of His-GFP-MTLS on MT tips compared to MT shafts (Figures 6A), which leads to an uneven distribution of the kinetic parameters along the MT. Our experimental data indicated that the formation of a TAC complex increased the probability of tubulation events and suppressed membrane detachment by promoting membrane association with MT tips (Figures 4H). TAC formation can be regarded as the assembly of an extra EB3-dependent adhesion domain at the MT tip, due to the enhancing of His-GFP-MTLS at the MT tip k_on-m-tip_ that we estimated is 2.5*k_on-m_ (Table 1, parameter 5) (Figures 6A-C). In the analysis of the *in silico* experiments, simultaneous movements of the membrane and MT tips were classified as TAC events, whereas other membrane spreading events were classified as sliding. Membrane tube retraction occurring independently of the MT tip was scored as a detachment event (Figures 6D and 4G). Similar to the experimental data (Figure 4H), the probability of successful sliding events observed in the simulations did not depend on the presence of EB3 (Figure 6E).

**Figure 6.**
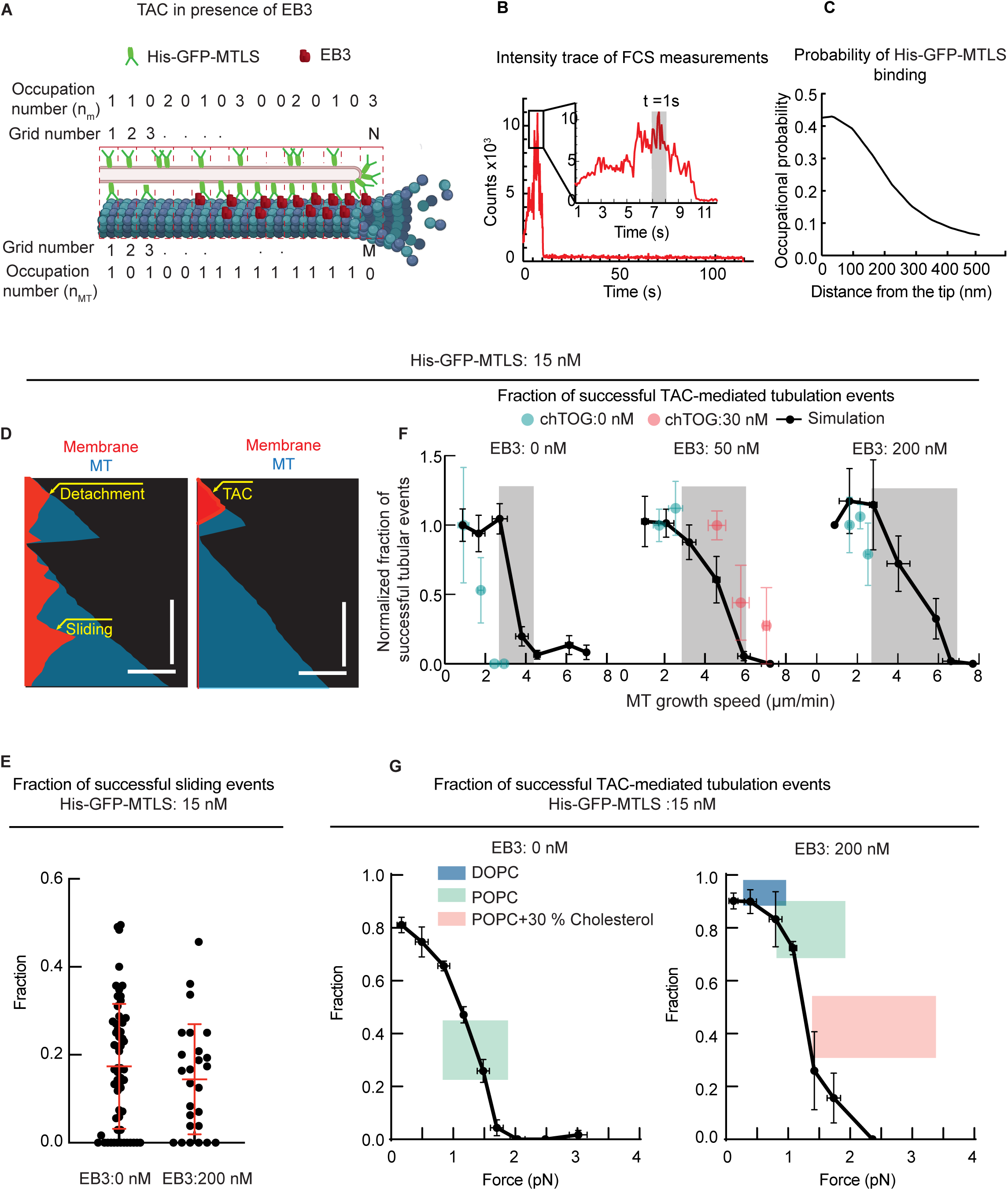
Simulations of TAC-based membrane tubulation and comparison of *in silico* and experimental data. (A) Kinetic schemes for the simulations of a membrane sliding at the MT tip in the presence of EB3. (B) Representative intensity time trace of a MT tip labeled with EB3-GFP recorded by FCS. The inset shows maximum intensity during a time window of 12 s. The grey rectangle marks the intensity in 1 s. (C) Estimated occupational probability of His-GFP-MTLS at the MT tip in the presence of 200 nM EB3. (D) Kymographs obtained from the stochastic simulations of membrane spreading in the presence of dynamic MTs. (E) Fraction of successful sliding tubulation events, calculated from simulations in the same way as in Figures 4H. EB3 0 nM, n=58 independent simulations (each represented by a black dot) and 1007 individual traces; EB3 200 nM, n=28 independent simulations and 348 individual traces. Error bars indicate SD. (F) Fraction of membrane deformations becoming tubular as a function of MT growth rate for simulation and experiments. Data were normalized to the average fraction obtained at the lowest MT growth rate. The grey rectangles mark the areas where the probability decays with the speed. 0 nM EB3, n=50 events, 7 experiments; 200 nM EB3, n=68 events, 9 experiments; 50 nM EB3, n=28 events, 7 experiments without chTOG; n=22 events, 4 experiments at 30 nM of chTOG. Error bars indicate SEM. (G) Fraction of membrane deformations becoming tubular as a function of the tube force. EB3 0 nM, n=59 independent simulations and 824 individual traces; EB3 200 nM, n=31 independents simulations and 424 individual traces. Error bars indicate SEM. The experimentally measured values are represented by rectangles due to the experimental uncertainty.

The analytical model predicted that there was a certain maximum speed at which a membrane could slide along a MT (Figure 5B and Equation 11 in Methods). Therefore, we expected that the TAC-driven membrane tubulation could be impeded at high MT growth rates. To test this hypothesis, we performed simulations and quantified the fraction of TAC events at different MT growth rates (Figure 6F). The simulations predicted that the fraction of successful tubular deformations was approximately constant when MTs grew with a speed of 1-3 µm/min (Figure 6F). When MT growth rates were increased, the system approached a transition regime where the probability of tubular deformations rapidly decayed, indicating that the formation of a TAC complex was limited by the maximum sliding speed of the membrane. The inclusion of EB3 in the simulation increased the fraction of TAC-mediated membrane tubulation at higher MT growth rates, because of a higher *k_on_* and thus higher maximum spreading speed (Figure 6F).

We next investigated the match between our simulations and the experimental data, focusing on the TAC events. In the absence of EB3, the transition to low tubulation probabilities was found at lower MT growth speeds than in the presence of EB3, indicating that EB3 enhanced the capacity of the membrane to follow the MT tip, in line with the predictions (Figure 6F). We note that the *in silico* results matched the experimental data in the presence of EB3 better than in the absence of EB3 (Figure 6F).

Since MT growth speeds in the presence of EB3 were always below the range where, according to the simulations, the transition to low tubulation probability occurred, we aimed to increase the MT growth speed in our assays. To achieve this, we added the human MT polymerase chTOG, since it has been shown that a combination of the frog homologue of chTOG, XMAP215, and the EB3 homologue EB1 strongly increased MT growth rate *in vitro* (Zanic et al., 2013). In the presence of chTOG, the MT polymerization rate was indeed increased, whereas membrane tubulation probability was reduced, just as our simulations predicted (Figure 6F). Taken together, these results support the idea that TAC formation is limited by the MT growth rate.

### Estimation of forces sustained by TACs during membrane spreading

We next combined simulations and experiments to estimate the forces that TACs could sustain. To do this, we performed stochastic simulations at different values of the force needed to pull a membrane tube (Derényi et al., 2002). We then estimated the probability of TAC formation in GUVs with different elastic properties and plotted it as the function of the force (Figure 6G). To further compare our simulations with experiments, we prepared GUVs with different values of the bending modulus by adding cholesterol, which makes membranes more stiff (Arriaga et al., 2017; Rodriguez-Garcia et al., 2009; Rodríguez-García et al., 2011). As expected (Campàs et al., 2008; Derényi et al., 2002; Koster et al., 2003), when the force required to make a tube increased, the probability of tubulation decreased. The estimated maximal force above which no tubes could be pulled by spreading was ∼ 1.5-2 pN, depending on the EB3 concentration, with the fraction of successful tubulation events at these forces being ∼5%. At the same lipid composition and thus the same membrane rigidity, the presence of EB3, which promotes TAC assembly, increased the chances of pulling a tube both in the simulations and in the experiments (Figure 6G). Altogether, our simulations, which are based on experimentally measured parameters and incorporate known physical effects related to membrane mechanics, including elastic membrane properties, provide a good quantitative explanation of the experimentally observed features of our *in vitro* reconstitution system.

### Measurement of TAC-mediated force production

Since direct measurement of the force acting on a spreading membrane was challenging, we turned to other artificial cargos. First, we asked whether a quantum dot (Qdot) could be transported by adhesive interactions with growing MT tips without external load. We added streptavidin-coated Qdots in the presence of EB3 and a biotinylated version of the dimeric MTLS (Bio-mCherry-MTLS) to a TIRFM assay in conditions previously identified to promote MT-driven membrane spreading (Figure 7A). When both EB3 and Bio-GFP-MTLS were present, we observed Qdots travelling steadily with the growing MT tips (Figure 7B, Video S7). EB3 alone, without Bio-mCherry-MTLS, did not promote Qdot binding to MTs, while Bio-mCherry-MTLS alone promoted Qdot binding only to the MT shafts, but not to the tips (Figure 7B). These results demonstrate that although EB proteins and their binding partners accumulate at MT ends by diffusion and exchange rapidly at MT tips(Bieling et al., 2007; Dragestein et al., 2008), a cluster of EB ligands can undergo processive EB-dependent motility on growing MT ends.

**Figure 7.**
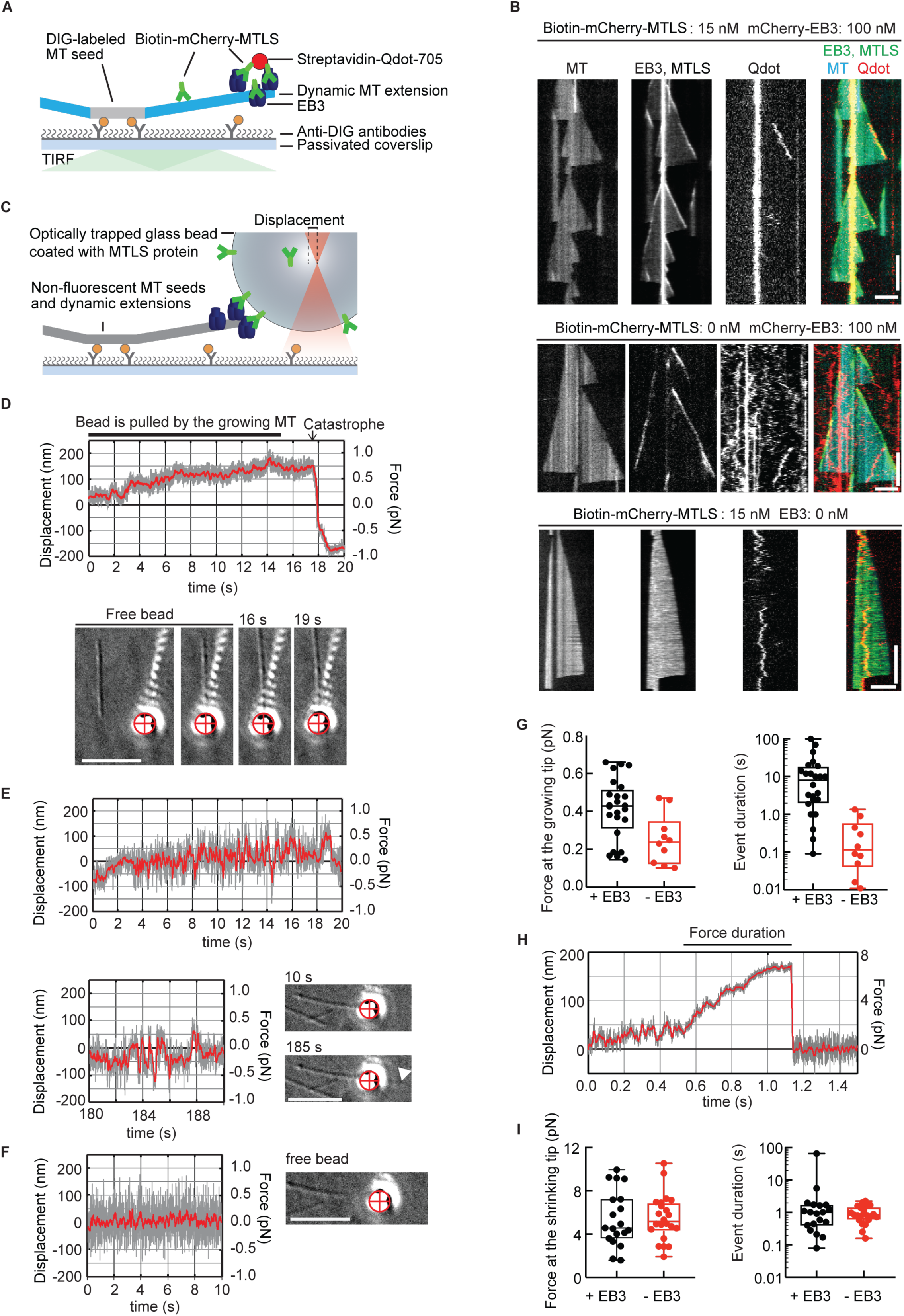
Cargo transport and force generation by growing and shrinking MT tips. (A) Schematics of the TIRFM experiment setup. (B) Kymographs showing MTs polymerized from HiLyte-488 tubulin (cyan), bio-mCherry-MTLS and mCherry-EB3 (green) and Qdot705-streptavidin (red). Notation above the kymographs describes the experimental conditions. Scale bars: horizontal (5 μm), vertical (60 s). (C) Setup of the optical trap experiments. (D) An example of a QPD trace in the presence of EB3 showing the raw signal at 1 kHz (grey) and the signal after smoothing with a running average of 100 points (red). Images at the bottom show corresponding frames from the DIC video (Video S8). The red cross marks the centre of the optical trap. Scale bar: 5 μm. (E) Analogous experiment in the absence of EB3. Bottom trace: the fast back and forth movements continued after the MT tip grew past the position of the trapped bead (white arrowhead). Scale bars: 5 μm. (F) Example trace and image of a free, unattached bead in a trap. Scale bar: 5 μm. (G) Force amplitude (left) measured at the growing MT tip, and force duration (right) in the presence (black, n = 24) or absence (red, n = 10) of 100 nM EB3. Horizontal line: median; box: 25-75%, whiskers: minimal and maximal values. (H) Example trace of an MTLS-coated bead in the absence of EB3 pulled by a shortening MT tip showing the raw signal at 10 kHz (grey) and the signal after smoothing with a running average of 100 points (red). (I) Force amplitude (left) measured at the shrinking MT tip, and duration (right) in the presence (black, n = 19) or absence (red, n = 22) of 100 nM EB3.

Next, we adapted the assay for direct measurement of the force sustained by TACs at growing MT ends. Neutravidin-coated glass beads were incubated with Bio-mCherry-MTLS and added to a chamber containing dynamic MTs, EB3 and Bio-mCherry-MTLS. A bead was optically trapped using low-power laser tweezers (stiffness ∼0.005 pN/nm) and placed in front of a growing MT tip (Figure 7C). Upon binding to a MT, the bead was initially pulled away from the trap center, and then gradually returned to zero position under assisting load applied by the optical trap (Video S8). We then used differential interference contrast (DIC) microscopy and high-resolution read-out from the quadrant photo detector (QPD) to ask whether the MT-bound bead continued its motion with the growing MT tip past the zero-force position against the opposing load applied by the optical trap. We found that when both EB3 and MTLS were present, the beads were displaced by growing MT tips past the zero position of the trap for seconds (Figure 7D). This displacement lasted until a MT catastrophe, when the beads, which presumably retained their association with the MT tip, were pulled out of the trap by the force generated by MT shortening (Figure 7D).

In the absence of EB3, the beads returned to zero-force position but did not continue to move steadily with the growing MT tip. Instead, we observed frequent and brief bead motions back and forth along the MT which continued when the MT tip grew past the position of the bead (Figure 7E). These motions were clearly different from normal fluctuations of a trapped bead that was free from any interactions with MTs or glass (Figure 7F). To quantify the effect of EB3 on the QPD signal, we measured the duration and the amplitude of the first positive bead displacement after the initial return to the zero position (Figure 7G). Only forces above 0.1 pN (the resolution of our instrument) were quantified. We found that most events in the absence of EB3 were shorter than 1 s, while in the presence of EB3 they lasted for 15 ± 5 s (mean ± SEM, N = 24). The forces generated when both EB3 and Bio-mCherry-MTLS were present were in the range of 0.1-0.7 pN, while the forces observed without EB3 were significantly lower. Taken together, our data indicate that TAC formation allows growing MTs to exert adhesion-based forces in the range of 0.5 pN.

To measure the forces that are exerted by dTACs during MT shortening, we performed the same experiment using a much stiffer trap (0.03-0.05 pN/nm). In these conditions the forces in the direction of MT growth were barely distinguishable against the bead’s thermal fluctuations. However, forces in the direction of MT shortening did not result in the beads escaping the trap, as was observed for experiments with the soft trap (Figure 7D, Video S8). Instead, the beads were pulled from the center of the optical trap by the shortening MT end, followed by detachment (Figure 7H). We observed forces in the range of 2-10 pN that typically lasted for 0.2-2 s, independent of the presence or absence of EB3 (Figure 7I). We have thus reconstituted EB3-dependent force coupling of TACs and EB-independent force coupling of dTACs.

## Discussion

In this study, we reconstituted MT-based membrane tubulation using a minimal system sufficient to drive formation of long membrane tubes by three mechanisms: sliding, TAC and dTAC, all of which were previously described in cells (Grigoriev et al., 2008; Guo et al.; Waterman-Storer et al., 1995; Waterman-Storer and Salmon, 1998). Previously, the sliding mechanism was attributed to motor proteins (Grigoriev et al., 2008; Waterman-Storer and Salmon, 1998; Woźniak et al., 2009). However, our data show that *in vitro*, membranes can spread on MT shafts independently of motors due to the adhesion mediated by a protein that can connect a lipid bilayer to MTs. Also TAC and dTAC events can be driven by a single protein that can connect membranes to MTs, raising the question about the contribution of such biochemically simple mechanisms to shaping the ER and other membrane organelles in cells.

Although membrane tubes could form on dynamic MTs in the presence of a MT-binding protein alone, the efficiency of membrane tubulation was strongly increased in the presence of EB3, to which this protein could bind. Previous work in cells showed that the formation of theTAC complex requires the interaction between the MTLS-containing transmembrane ER protein STIM1 and EBs (Grigoriev et al., 2008), but it remained unclear whether additional molecules are needed for membrane tubulation by this mechanism. Here, we showed that a combination of an EB protein and its membrane-associated partner is sufficient to promote efficient membrane tube extension by growing MT ends. This mechanism relies on molecules that, unlike molecular motors, do not undergo processive motion, but rather concentrate at growing MT ends by a diffusion-based mechanism because of an increased affinity for the MT tip structure (Bieling et al., 2007; Dragestein et al., 2008). In spite of their fast turnover, clustering of MT tip-tracking proteins on a specific structure, such as a bead or a membrane tip, is sufficient to bias the displacement of this structure and thus induce its processive motility. Importantly, the EB3-dependent TAC mechanism was particularly important when MTs grew rapidly, and this likely explains its relevance in cells, where MTs grow with at a rate reaching 10-20 μm/min.

Furthermore, we observed that the presence of a protein with dual affinity to membranes and MT shafts was sufficient to pull membrane tubes by shrinking MTs (dTAC mechanism). Recent imaging work revealed dTAC-based ER remodeling in cells, and it was proposed that this mechanism employed factors different from those specific for TAC generation (Guo et al., 2018). However, our data show that the same molecules can in principle support both mechanisms. In fact, it is possible that any diffusible ER-resident protein with a strong affinity for MT lattices might be able to induce dTAC formation.

Our measurements indicated that in the conditions used, MT tip-localized EB3-MTLS complexes can mediate force generation in the range of ∼0.5 pN. This value is in line with a recent study, which proposed that the interaction of a MT tip-bound EB protein with kinesin-14 can reverse the movement direction of this kinesin by exerting a force with a magnitude below 1 pN (Molodtsov et al., 2016). Based on the minimal value of tension and the membrane bending modulus measured by flickering spectroscopy, a force in the range of ∼0.5 pN is sufficient to cause tubulation of membranes that are not under tension. However, based both on our simulations and experiments, the forces generated during TAC events when membranes were under some tension were estimated to be in the range of 1-2 pN. The discrepancy between this estimate and the force measurements at growing MT tips can be attributed to the fact that the additional force is likely sustained by membrane adhesion along the MT lattice, whereas this contribution is absent in the bead-based assay. Unfavorable geometry of bead-MT contacts could also reduce apparent force measured in the center of the bead compared to the MT-generated force acting at the bead surface (Volkov et al., 2013). Furthermore, the membrane-MT interface, with the membrane partly wrapping around the MT cylinder, might lead to a higher number of contacts that together can sustain higher forces compared to a MT interface with a spherical bead. The forces generated by the combination of membrane-MT shaft interactions and the TAC complex are of the same order of magnitude as the forces generating membrane tubes by one-dimensional wetting (Charles-Orszag et al., 2018). Interestingly, the forces generated when the same MT-membrane connecting molecules are attached to shrinking MTs are an order of magnitude higher than those mediated by TACs, possibly because the attachment geometry is significantly different at depolymerizing ends.

To conclude, our data show that the three different mechanisms of MT-driven membrane tubulation can be explained by the formation of transient bonds between MTs and membranes, and that a biased self-spreading of the lipid bilayer on MTs can account both for the sliding and TAC mechanisms. Whereas previous work has provided the proof of principle for membrane spreading on fibers, the employed systems did not aim to reconstitute molecular mechanisms that are operational in the cell: they used static actin filaments decorated with neutravidin and biotinylated membranes (Charles-Orszag et al., 2018), or a combination of anionic porphyrin J-aggregates with DOPC liposomes (Sugikawa et al., 2017). In contrast, the system we used recapitulates the dynamics and affinities found in naturally occurring MT-binding proteins and dynamic MTs, and therefore it is of direct relevance to the MT-membrane contacts found in cells. MT-based membrane tubulation plays a major role in shaping the ER (Westrate et al., 2015) and is also important for the morphogenesis of other organelles, such as mitochondria (Kanfer et al., 2017) and recycling endosomes (Delevoye et al., 2016). Since numerous non-motor MT-membrane linking proteins are known, they are likely to contribute to intracellular organelle organization through adhesion-based mechanisms described in this study.

## Methods

### Preparation of vesicles

1-palmitoyl-2-oleoyl-glycero-3-phosphocoline (POPC), 1,2-dioleoyl-sn-glycero-3-[(N(5-amino-1-carboxypentyl)iminodiacetic acid)succinyl](nickel salt) (DOGS-Ni-NTA), 1,2-dioleoyl-sn-glycero-3-phosphoethanolamine-N-(lissamine rhodamine B sulfonyl) (ammonium salt) (Rh-PE), 1,2-dioleolyl-sn-glycero-3-phosphocoline (DOPC) and 1-oleoyl-2-{6-[(7-nitro-2-1,3-benzoxadiazol-4-yl)amino]hexanoyl}-sn-glycero-3-phosphocoline (NBD-PC) were purchased from Avanti Polar Lipids. Cholesterol was purchased from Sigma Aldrich. The lipid mixtures were composed of 94.95% POPC, 5% DOGS-Ni-NTA, 0.05% (Rh-PE/NBD-PC) or 94.95% DOPC, 5% DOGS-Ni-NTA, 0.05%Rh-PE. For experiments with cholesterol, the lipid mixture was 64.95% POPC, 5% DOGS-Ni-NTA, 0.05% Rh-PE and 30% Cholesterol (expressed as molar proportions). GUVs were prepared by electroformation (Mathivet et al., 1996) on two conductor indium tin oxide (ITO)-coated glass slides (Sigma Aldrich). Briefly, ten microliters of a lipid suspension (0.1 mg /mL) in chloroform were spread over the conductor surface. A chamber was made with non-adhesive silicone spacer (0.8 mm depth, Sigma Aldrich); after solvent evaporation the films were hydrated with sucrose solution (300 mM), and the electrodes were connected to an AC power supply (1V, 10 Hz) for 3.5 hours and (1.5V, 5 Hz) for 30 min to ensure good detachment of the GUVs from the ITO glass slides.

### Protein purification

His-mCherry-EB3 was purified from *E. coli* as described previously(Montenegro Gouveia et al., 2010). mCherry-EB3 was cloned into a pET based bacterial expression vector with a N-terminal 6xHis tag followed by a 3C cleavage site using the restriction free positive selection method (Olieric et al., 2010). Protein production was performed in the *E. coli* expression strain Bl21(DE3) in LB broth by inducing with 0.5 mM IPTG at an OD_600_ of 0.4 to 0.6 overnight at 20 °C. Cells were harvested by centrifugation at 4 °C, 3500 x g for 15 min and lysed by sonication in a buffer containing 50 mM HEPES, pH 8.0, 500 mM NaCl, 10 mM Imidazole, 10% Glycerol, 2 mM β-mercaptoethanol, and proteases inhibitors (Roche). The crude extracts were cleared by centrifugation at 20,000 x g for 20 min and the supernatants were filtered through a 0.4 micron filter before purification.

Protein purification was performed by immobilized metal-affinity chromatography (IMAC) on HisTrap HP Ni^2+^ Sepharose columns (GE Healthcare) at 4°C according to the manufacturer’s instructions. The 6xHis tag was cleaved overnight by 3C protease during dialysis against lysis buffer (without proteases inhibitors). The cleaved sample was reapplied onto the IMAC column to separate the cleaved product from its tag and potentially uncleaved protein. Processed protein was concentrated and further purified on a HiLoad Superdex 200 16/60 size exclusion chromatography column (GE Healthcare) equilibrated in 20 mM Tris HCl, pH 7.5, 150 mM NaCl, 2 mM DTT. Protein fractions were analyzed by Coomasie stained SDS-PAGE. Fractions containing mCherry-EB3 were pooled and concentrated by ultrafiltration. Protein concentration was estimated by UV at 280 nm and the pure mCherry-EB3 was aliquoted, flash frozen in liquid nitrogen and stored at −80°C.

His-GFP-MTLS corresponds to the previously described construct, which includes EGFP, the two-stranded leucine zipper (LZ) coiled-coil domain of GCN4 and the C-terminal 43 residues of human MACF2 (MACF43)(Honnappa et al., 2009), which was cloned between *Nde*I and *Bam*HI sites of pET28a (Novagen, Merck Millipore, Billerica, MA, USA) and purified as described previously (Buey et al., 2012).

Bio-mCherry-MTLS was assembled by PCR amplification from mCherry with an N-terminal biotinylation tag (the peptide MASGLNDIFEAQKIEWHEGGG, which serves as a substrate for the bacterial biotin-protein ligase BirA (de Boer et al., 2003)), LZ and MACF43. Bio-mCherry-MTLS was cloned into pET28a vector linearized with *EcoRI* and *SalI* restriction enzymes. Protein purification was performed from BL21 DE3 cells. Expression was induced at OD 0.6 with 1 mM IPTG and continued overnight at 20°C. After induction, bacteria were harvested by centrifugation. The supernatant was discarded and bacteria were resuspended in 5 ml/gram lysis buffer (50 mM Tris-HCl, pH 7.4, 300 mM NaCl, 1 mM dithiolthreitol (DTT), 0.5% Triton-X, Complete Protease Inhibitor and 0.2 mg/ml lysozyme.) After ∼15 min incubation, cells were sonicated 5 times for 30 seconds with intervals of 1 min. The extract was pre-cleared by centrifugation at 20.000 x g for 45 minutes and the supernatant was incubated for 1 hr with Streptactin Sepharose beads (GE Healthcare) which had been washed 3 times in the lysis buffer. Next, beads were washed with a solution containing 2.5 mM D-biotin in the lysis buffer and spun down. The supernatant then incubated with Ni-NTA beads (Qiagen) for 1 hr. Subsequently, the beads were again spun down and washed with the elution buffer containing 50 mM Tris-HCl pH 7.4, 150 mM NaCl, 300 mM imidazole, 1 mM DTT, 0.1% Triton. Finally, a PD-10 desalting column (GE Healthcare) and a concentrator (Vivaspin MWCO 10K, Satorius) were used to perform buffer exchange. The protein was snap-frozen with 10% glycerol.

Human ch-TOG was purified from HEK293T cells using the Strep(II)-streptactin affinity purification. ch-TOG expression construct was based on a previously described construct(Gutierrez-Caballero et al., 2015) that was a gift of S. Royle (University of Warwick, UK). Ch-TOG was cloned into a modified pEGFP-N1 vector with a StrepII tag. HEK 293T cells were cultured in DMEM/F10 (1:1 ratio, Lonza, Basel, Switzerland) supplemented with 10% FCS, transfected using polyethylenimine (PEI, Polysciences) and harvested 2 days after transfection. Cells from a 15 cm dish were lysed in 500 µl of lysis buffer (50 mM HEPES, 300 mM NaCl and 0.5% Triton X-100, pH 7.4) supplemented with protease inhibitors (Roche) on ice for 15 minutes. The supernatant obtained from the cell lysate after centrifugation at 21,000 x g for 20 minutes was incubated with 40 µl of StrepTactin Sepharose beads (GE) for 45 minutes. The beads were washed 3 times in the lysis buffer without the protease inhibitors. The protein was eluted with 40 µl of elution buffer (50 mM HEPES, 150 mM NaCl, 1 mM MgCl_2_, 1 mM EGTA, 1 mM dithiothreitol (DTT), 2.5 mM d-Desthiobiotin and 0.05% Triton X-100, pH 7.4). Purified proteins were snap-frozen and stored at −80 °C.

### In vitro reconstitution assays

*In vitro* assays with dynamic MTs were performed as described previously(Mohan et al., 2013). First double cycled GMPCPP MT seeds were prepared by incubating tubulin mix containing 70% unlabeled porcine brain tubulin (Cytoskeleton), 18 % biotin-tubulin (Cytoskeleton) and 12 % rhodamine-tubulin (Cytoskeleton) at a total tubulin concentration of 20 µM with 1 mM GMPCPP (Jena Biosciences) at 37° C for 30 minutes. MTs were pelleted by centrifugation in an Airfuge for 5 minutes and then depolymerized on ice for 20 minutes. After this step, a second round of polymerization at 37° C with 1mM GMPCPP was performed. MT seeds were pelleted as above and diluted 10 fold in MRB80 buffer (80 mM piperazine-N,N[prime]-bis(2-ethanesulfonic acid, pH 6.8, supplemented with 4 mM MgCl_2_, and 1 mM EGTA) containing 10% glycerol, snap frozen in liquid nitrogen and stored at −80° C.

Flow chambers were assembled by sticking plasma-cleaned glass coverslips onto microscopic slides with a double sided tape. The chambers were treated with 0.2 mg/mL of PLL-PEG-biotin (Surface Solutions, Switzerland) in MRB80. After washing with the assay buffer MRB80, the chambers were incubated with 1 mg/mL NeutrAvidin (Invitrogen). Then 2-3 µL of MTs seeds were diluted in 80 µL of glucose 320 mM and attached to the biotin-NeutrAvidin links. The reaction mixtures supplemented with 20 µM of tubulin, 50 mM KCL,0.1% Methyl cellulose,0.5 mg/mL k-casein, 1 mM GTP, an oxygen scavenging system (200 μg/mL catalase, 400 μg/mL glucose oxidase, 4mM DTT), and 100 nM of SiR-Tubulin (Cytoskeleton) was added to the flow chamber after centrifugation in an Airfuge for 5 minutes at 119,000 g. In the experiments with GUVs, the vesicles were added after centrifugation. The flow chamber was sealed with vacuum grease, and dynamic MTs with and without GUVs were imaged at 30°C using TIRF microscopy.

### TIRF microscopy

TIRF microscopy was performed on an inverted microscope Nikon Eclipse Ti-E (Nikon) with a perfect focus system. The setup was equipped with a Nikon CFI Apo TIRF 100×1.49 N.A oil objective, Photometrics Evolve 512 EMCCD (Roper Scientific) or CoolSNAP HQ2 CCD camera (Roper Scientific) and controlled with the Metamorph 7.7.5 software. The final magnification was 0.063 μm/pixel, which includes the 2.5x magnification introduced by an additional lens (VM lens C-2.5x, Nikon). The temperature was controlled by a stage top incubator INUBG2E-ZILCS (Tokai Hit). The microscope was equipped with TIRF-E motorized TIRF illuminator modified by Roper Scientific France/PICT-IBiSa, Institut Curie. For excitation lasers we used 491 nm 100 mW Stradus (Vortan), 561 nm 100 mW Jive (Cobolt) and 642 nm 110 mW Stradus (Vortran). We used an ET-GFP 49002 filter set (Chroma) for imaging proteins tagged with GFP and NBD-PC lipid, an ET-mCherry 49008 filter set (Chroma) for imaging X-Rhodamine labelled tubulin, Rh-PE lipid or mCherry-EB3 and an ET-405/488/561/647 for imaging SiR tubulin labelled MTs or Hilyte 647 porcine brain tubulin. We used sequential acquisition for triple-color imaging experiments.

### Spinning disk microscopy

Spinning disk confocal microscopy was performed on a Nikon Eclipse Ti microscope equipped with a perfect focus system (Nikon), a spinning disk-based confocal scanner unit (CSU-X1-A1, Yokogawa, Japan), an Evolve 512 EMCCD camera (Roper Scientific, Trenton, NJ) attached to a 2.0X intermediate lens (Edmund Optics, Barrington, NJ), a super high pressure mercury lamp (C-SHG1, Nikon), a Roper scientific custom-ordered illuminator (Nikon, MEY10021) including 405 nm (100 mW, Vortran), 491 nm (100 mW, Cobolt) 561 nm (100 mW, Cobolt) and 647 nm (100 mW, Cobolt) excitation lasers, a set of BFP, GFP, RFP and FarRed emission filters (Chroma, Bellows Falls, VT) and a motorized stage MS-2000-XYZ with Piezo Top Plate (ASI). The microscope setup was controlled by MetaMorph 7.7.5 software. Images were acquired using Plan Fluor Apo VC 60x NA 1.4 oil objective. The 3D image reconstruction was carried out using Huygens Professional version 18.04 (Scientific Volume Imaging, The Netherlands).

### Quantification of His-GFP-MTLS intensity along MTs

To build the average distribution of His-GFP-MTLS intensity at the MT tip (Figure S1B,D), we generated mean intensity profiles of 6 pixel (400 nm) thick lines with a length 2-3 µm along MTs, with the middle point positioned approximately at the MT tip (Aher et al., 2018; Demchouk et al., 2011). Previously (Aher et al., 2018), we extracted the fluorescence profile along a MT and fitted it with the error function, which determined MT tip position. Since the profile of His-GFP-MTLS intensity had a shape of a comet, fitting it to the error function was not possible (Figure S1E). In order to use the error function to determine the position of MT tip, the maximum intensity from the His-GFP-MTLS protein fluorescence profile was assigned to all preceding points along the MT(Figure S1E)(Seetapun et al., 2012). Then as described previously(Aher et al., 2018; Demchouk et al., 2011), the intensity profiles *I(x)* were fitted with the error function shifted in *x* as:

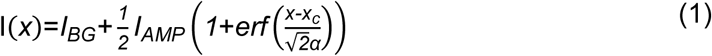

where *I_BG_* corresponds to the intensity of background, *I_AMP_* the amplitude of the fluorescent signal, *x_c_* is the position of the MT tip and *α* the degree of tip tapering convolved with the microscope’s point spread function. Each profile was shifted by its *x_c_* value and background was subtracted, the 0 distance was defined as the position of the MT tip. To generate the binding curve of the His-GFP-MTLS protein (Figure S1C), profile intensities were measured at different EB3 concentrations on a single day with the same microscopy settings (Maurer et al., 2011). To estimate the mean His-GFP-MTLS intensity along the MTs (Figure 3B), we generated mean intensity profiles of 6 pixel (400 nm) thick lines along whole MTs.

### Analysis of the shape of tubular membrane network

To analyze the shape of a tubular network, we first generated a single image containing the maximum intensity projection of the time lapse movie using Fiji (Figure S2)(Schindelin et al., 2012). A custom MATLAB script was employed to segment the network using a supervised segmentation pipeline. Segmented images (Figure S2) were used to measure the perimeter and the area using built-in MATLAB functions from the image processing toolbox. Sphericity was calculated by dividing the total area by the squared value of the perimeter(Legland et al., 2016). The sphericity was normalized as:

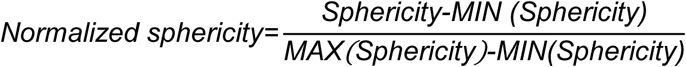

where *max(sphericity)* and *min(sphericity)* were defined as the maximal and minimal value of the experimentally measured sphericity, respectively. Original segmented images were converted into nodes and branches (skeletonization) using built-in MATLAB functions. Skeleton representations of the networks were used to measure the total length of membrane tubes as the sum of the lengths of all detected branches (Figure S2).

### Analysis of thermal membrane fluctuations

Analysis of thermal fluctuations was performed as described previously (Pecreaux et al., 2004; Rodriguez-Garcia et al., 2009; Rodriguez-Garcia et al., 2015). GUVs were visualized using a phase contrast microscope (Nikon TE2000) with a 60x NA 1.4 oil objective. Fluctuations of the vesicle radius were measured at the equatorial plane by videomicroscopy. We acquired 1000 time lapse images of fluctuating GUVs. Each vesicle profile was digitalized and the GUV-equatorial fluctuations, namely *δR(x,t)=R_0_-r_e_(x,t)* were measured as local deviations of the equatorial radius *r_e_* at each point of the GUV contour from an average value, *R_0_*, where *R_0_* is calculated with respect to the center of mass of the GUV profile. The equatorial fluctuations were expanded in a series of discrete Fourier modes *n* as *δR(x,t)/R_0_=Σ_q_*(*a_q_sin*(*qx*)+*b_q_cos*(*qx*)), where *a_q_* and *b_q_* are the Fourier coefficients and *q*=*n/R_0_* (with n=2,3,…50). The series were truncated at *n =* 50.

The spectrum of the membrane fluctuations in the equatorial plane *P_eq_* (in units of length^^3^) by calculating time-averages of quadratic fluctuation amplitudes: *P_eq_*=*A*\*(〈|*c_n_*|^2^〉-〈|*c*_n_|〉^2^), where 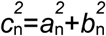 and *A=*〈π*R_0_*〉*^3^/2* (Figure S3, left). Since the amplitudes of the fluctuations estimated from experiments depended on the GUV mean radius 〈*R_0_*〉, non-dimensional averaged curves were obtained after dividing each individual spectrum by *A=*〈π*R_0_*〉*^3^/2* (Figures 3G and S3 right)(Pecreaux et al., 2004; Rodriguez-Garcia et al., 2009). The estimated amplitude of the Fourier modes included errors due to the finite spatial resolution of the images (pixelization). This noise introduces in the fluctuations a systematic component, which could become dominant at high fluctuation modes (Gittes et al., 1993; Pecreaux et al., 2004). Consequently, the experimentally measured amplitudes were corrected by subtracting the noise as *P_eq_* (corrected) = *P_eq_* − [(π〈*R_0_*〉*^3^/2*)*var(*c_n_*)], where var(*c_n_*) represents the variance of *c_n_* as explained in (Pecreaux et al., 2004). A detailed derivation of the formulas for error propagation is provided in the Appendix C of (Pecreaux et al., 2004). In addition, we quantified the minimal fluctuation that we can estimate by measuring the fluctuation spectrum of a fixed object in the focal plane (Figure 3G). As a model for a fixed object, we used a transparent circular disc of radius 10 μm, printed in a Quartz photomask coated with chrome (Toppan). Every fluctuation amplitude below the estimated resolution limit was discarded during fitting. A correction to account for to the finite integration time of the camera (Méléard et al., 1992; Pecreaux et al., 2004) was also included in the analysis.

Mechanical parameters of the GUVs were obtained by fitting the experimental mode amplitudes to the theoretical spectrum for a planar membrane (Pecreaux et al., 2004; Rodriguez-Garcia et al., 2009). Detection of the vesicle contours, post-processing and fitting procedures were carried out using a custom MATLAB script as described previously (Rodriguez-Garcia et al., 2009).

### Analysis of MT and membrane dynamics

To analyze the dynamics of MTs and membrane tubes, kymographs were generated using Fiji (KymoResliceWide plugin, https://github.com/ekatrukha/KymoResliceWide). MT catastrophe frequency was determined as the ratio of the total number of catastrophes observed for a particular MT to the total time a MT spent growing. MT growth rate, as well as the speed of growing membrane tubes were determined manually from kymographs. To estimate the probability of tubular deformations and detachment events, we selected MTs growing beyond the visible contour of the GUV (Figure 4G) and divided the number of observed tubular membrane deformations by the total number of MT growth events. To estimate the probability that a deformation stopped because the membrane tip detached from the MT tip, we divided the total number of detachment events by the total number of successful membrane deformations (Figure 4G,J).

To determine the spreading speed of a membrane tube, time lapse images of GUVs interacting with MTs were recorded at 3 s intervals with 200 ms exposure time for 5 or 10 min. Intensity profiles of membrane tubes labeled with Rh-PE were collected at different time points by averaging across 6-pixel (400 nm) wide lines. The position of a tube end was determined by fitting every profile with the error function using a custom written MATLAB script (Demchouk et al., 2011) (Figure S5A). Membrane tube spreading speed was calculated by differentiating the vector of the tube end position over time.

### Reconstitution of Qdot transport with MT tips

Digoxigenin (DIG)-labelled tubulin was prepared as described previously (Hyman, 1991). All other tubulin products were purchased from Cytoskeleton. Chambers were prepared and imaged as described previously (Volkov et al., 2018). In brief, silanized slides and coverslips were assembled into chambers using double-sided tape and functionalized with anti-DIG IgG (Roche), then passivated with 1% Tween-20. Double-cycled, DIG-labelled GMPCPP-stabilized seeds were introduced, followed by the reaction mix containing 80 mM K-Pipes pH 6.9, 50 mM KCl, 1 mM EGTA, 4 mM MgCl_2_, 1 mM GTP, 0.1% methylcellulose, 1 mg/ml κ-casein, 4 mM DTT, 0.2 mg/ml catalase, 0.4 mg/ml glucose oxidase, 20 mM glucose, 20 μM tubulin (5% labelled with HiLyte-488) and 30 pM Qdot705-streptavidin (Thermo Fischer) with or without the addition of 15 nM bio-mCherry-MTLS and 100 nM mCherry-EB3.

Images were acquired using Nikon Ti-E microscope (Nikon, Japan) with the perfect focus system (Nikon) equipped with a Plan Apo 100X 1.45 NA TIRF oil-immersion objective (Nikon), iLas^2^ ring TIRF module (Roper Scientific) and a Evolve 512 EMCCD camera (Roper Scientific, Germany). The sample was illuminated with 488 nm (150 mW for HiLyte488-tubulin), 561 nm (100 mW for mCherry-tagged proteins) and 642 nm (110 mW, for Qdot-705) lasers through a quad-band filter set containing a ZT405/488/561/640rpc dichroic mirror and a ZET405/488/561/640m emission filter (Chroma). Images were acquired sequentially with MetaMorph 7.8 software (Molecular Devices, San Jose, CA). The final resolution was 0.16 µm/pixel. The objective was heated to 34°C by a custom-made collar coupled with a thermostat, resulting in the flow chamber being heated to 30°C.

### Optical trapping

1 μm glass beads functionalized with carboxy groups (Bangs Laboratories) were conjugated with PLL-PEG (Poly-L-lysine (20 kDa) grafted with polyethyleneglycole (2 kDa), SuSoS AG, Switzerland) supplemented with 10% v/v of PLL-PEG-biotin as described(Volkov et al., 2018). The bead surface was then saturated with Neutravidin and then Bio-mCherry-MTLS. Flow chambers were prepared as described above. Reaction mix contained 80 mM K-Pipes pH 6.9, 50 mM KCl, 1 mM EGTA, 4 mM MgCl_2_, 1mM GTP, 1 mg/ml κ-casein, 4 mM DTT, 0.2 mg/ml catalase, 0.4 mg/ml glucose oxidase, 20 mM glucose and either 25 μM unlabelled tubulin with addition of 100 nM mCherry-EB3, or 10 μM unlabelled tubulin in the absence of EB3. Experiments were carried out at 25°C.

DIC microscopy and optical trapping were performed as described previously (Volkov et al., 2018). Measurements were performed at nominal trap power of 0.4W which resulted in a typical stiffness of 0.03-0.05 pN/nm. QPD signal was recorded at 10 kHz. For experiments measuring forces in the direction of the MT growth, which were expected to be in the sub-pN range, the 0.2W trap (the lowest setting for our instrument) was additionally softened down to 4-6 ·10^-3^ pN/nm using a circular polarizing filter placed in the wave path of the trapping laser. QPD signal was recorded at 10 kHz and down-sampled to 1 kHz for analysis.

### Single molecule measurements

His-GFP-MTLS single molecule binding events to MTs were recorded using TIRF microscopy (Figure S5C) as described previously (Sharma et al., 2016). Briefly, the assay was performed at 0.12 nM concentration of His-GFP-MTLS with high speed acquisition (50 ms/frame). 200 nM mCherry-EB3 was added as a tracer to detect MT plus ends. The number and dwell times of the binding events were extracted from kymographs that were generated using Fiji (KymoResliceWide plugin). The distributions of the dwell times were fitted with an exponential function, giving the dissociation rates from MT lattice and from MT tips (Figure S5D,E). The binding constant rate was determined by measuring the number of single molecule binding events per MT lattice length and assuming a length of 200 nm for the MT tip (Figure S5F).

The diffusion coefficient of His-GFP-MTLS on MT lattice *D_MT_* (Table 1, parameter 14) was derived from the analysis of the mean square displacement (MSD) of single molecule spots diffusing along MTs (Sharma et al., 2016)(Figure S5C,G). Coordinates of diffusing spots were obtained and linked across frames using the Fiji plugin TrackMate (Tinevez et al., 2017). MSD analysis was performed using MSDanalyzer MATLAB routine (Tarantino et al., 2014). 25% of each MSD curve excluding zero was used for fitting. To compute the mean *D_MT_* we only considered values where the coefficient of determination *R^2^* of the fitting was above 0.8 (n=95)(Figure S5G).

### Fluorescence correlation spectroscopy

We used z-scan Fluorescence Correlation Spectroscopy (z-FCS) to measure the diffusion coefficient of His-GFP-MTLS bound to GUVs (Benda et al., 2003). z-FCS measurements were performed with a Leica SP8 STED 3X microscope driven by LAS X and SymPhoTime (PicoQuant) software using a HC PLAPO 63x 1.2 NA water immersion objective. The sample was excited at 488 nm and time correlation was calculated at the time frequency of 600 kHz.

Confocal imaging was used to localize the top of the GUVs, starting from which a fast scan along the z-axis was performed. The position of the focus in z was set at 1 μm below or above of the plane with the maximum intensity. Afterwards, autocorrelation functions *G(t)* were measured at different positions along the z-axis in 0.2 μm steps. Three traces of 10 s duration were acquired at each position. The autocorrelation functions were fitted with the model considering one single species diffusing in 2 dimension described previously (Elson and Magde, 1974):

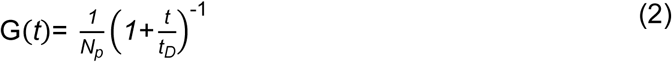

where *N_p_* is the average number of particles in the confocal volume, *t_D_* is the characteristic diffusion time and *t* is the lag time. The obtained average number of particles and diffusion times was plotted against the focus position *z* and fitted with the equations derived previously (Sorscher and Klein, 1980):

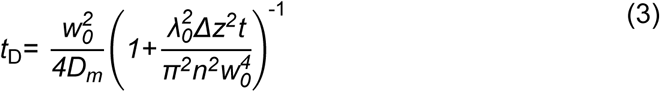

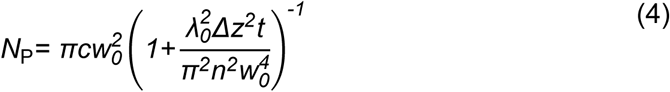

where *w_0_* (Table 1, parameter 18)is the radius of the beam in the focal plane, *D_m_* (Table 1, parameter 16) is the diffusion coefficient of the protein bound to membrane, *c* is the average concentration in the illuminated area, *n* is the refractive index of the medium and λ is the wavelength of the excitation light.

We used FCS to estimate the probability of finding His-GFP-MTLS bound to EB3-bound MT tip as a function of the distance from the MT tip. We reconstituted MT plus-end tracking of the His-GFP-MTLS (15 nM) and EB3 (200 nM) as before, but in the absence of GUVs. When we performed FCS experiments, we observed peaks in the fluorescence intensity trace. (Figure 6B). The peaks represented His-GFP-MTLS-labeled MT ends(Dragestein et al., 2008; Rodriguez-Garcia et al., 2018). From the intensity values next to the intensity peak we calculated the autocorrelation function as described above. The autocorrelation function *G*(*t*) was fitted using a triplet-state model for one fluorescent species (Widengren et al., 1994) as follows:

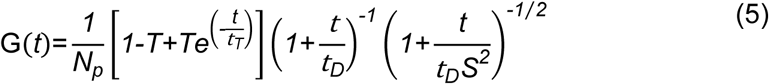

where *T* is the triplet decay fraction, *t_T_* is the lifetime of the triplet state, *t_D_* is the diffusion time of the fluorescent species and *S* is the length-to-diameter ratio of the focal volume set to 4. We then estimated the mean of the maximum numbers of His-GFP-MTLS molecules at the MT tip *<N_p-tip_*> as:

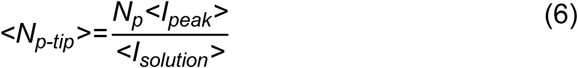

where <*I_peak_*> is the mean peak intensity measured during 1s (Figure 6B), and <*I_solution_*>is the averaged intensity measured after the peak (Figure 6B). It was assumed that the intensity ratio was proportional to the ratio of the MTLS copy number. Considering a growth rate for MTs of ∼ 2.8 μm/min we estimated that the maximum intensity recorded in 1 s corresponded to ∼46 nm of the MT tip (∼ 6 tubulin dimers). Since EB3 binds to MTs between protofilaments at the interface between four tubulin dimers but not at the seam (Maurer et al., 2012) we considered that within 6 dimer lengths of a 13 protofilament MT. Assuming that EB3 and MTLS interact at a ratio of 1:1, there were ∼ 60 potential binding sites for His-GFP-MTLS. Considering that at 200 nM EB3 all EB3 binding site could be occupied, we estimated that the maximum probability of finding His-GFP-MTLS bound to EB3-bound MT tip was <*N_p-tip_*>/60=28/60=0.46 (see Table1, parameter 13). Then we estimated the probability of His-GFP-MTLS binding along the MT length as the product of the maximum probability calculated above using normalized average His-GFP-MTLS intensity profiles measured by TIRF microscopy (Figure S1B).

### Estimation of the surface density of MTLS bound to the GUV surface in contact with MTs

To estimate the surface density of MTLS molecules bound to the GUV surface in contact with MTs (*ρ_MTLS-MT_*) (Table 1, parameter 21) we used FCS and fluorescence intensity analysis. As we described above we measured by FCS the average number of free MTLS molecules bound to the surface of the GUV in the confocal volume *N_p_* and the radius of the beam in the focal plane *w_0_*. Then we estimated the density of free MTLS molecules bound to the surface of the GUV (Table1, parameter 20) as the ratio between the average number of MTLS molecules and the area of the beam in the focal plane: *ρ_MTLS-MT_= N_p_* /*w ^2^*. Finally, the density of the MTLS molecules bound to the GUV surface in contact with MTs was determined as: *ρ_MTLS-MT_= ρ_MTLS_*(I_GUV-MTs contact_ /I_GUVS_)* where *I_GUV-MTs contact_* is the mean fluorescent intensity of the free MTLS bound to the surface of the GUV in contact with MTs *I_GUV-MTs_* and *I_GUVS_* is the fluorescent intensity of the free MTLS molecules bound to the surface of the GUV (Figure 3B).

### Physical modeling

#### Membrane sliding along MT lattice

To describe membrane sliding along MT lattice, a minimal one dimensional (1D) theoretical model was adapted from a previous publication (Boulbitch et al., 2001). We considered that the binding-unbinding reaction of His-GFP-MTLS to MT takes place in the close vicinity of the tip of a membrane deformation. The width of the reaction zone *d* in the 1D model was fixed to the length of one tubulin dimer, *d* = 8 nm. Moreover, we considered a reaction-dominated regime because of the high His-GFP-MTLS concentration at the GUV-MT interface (Figure 3A,B). For simplicity, we first considered only the tip of the tube that extended for a distance *L*. We assumed that the tip moves in a biased-random walk, stepping from one binding site to the next with forward rate *k_on-m_* and backward rate *k_off-m_*. The unbinding rate depends on the force barrier *F_m_(t)* to pull a membrane tube as described previously (Campàs et al., 2008).

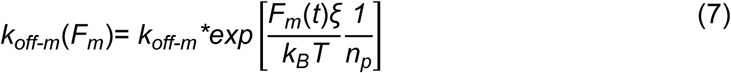

where *F_m_*(*t*)*=2π*(*2σ*(*t*)*κ*)^1/2^ (Derényi et al., 2002), *κ* is the bending modulus and *σ*(*t*) is the lateral tension of the membrane. The parameter *ξ* (Table 1, parameter 6) is the characteristic length of the potential barrier between bound and unbound states, *T* is the temperature, *k_B_* the Boltzmann’s constant and *n_p_* (Table 1, parameter 7) is the number of proteins bound at the tip. At the tip, the load is equally distributed between all the bonds due to the parallel organization (Figure 5A). The lateral tension of a flaccid membrane with initial tension *σ_0_*, increased during tube elongation as (Cuvelier et al., 2005):

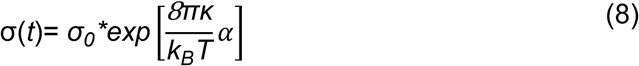

where *α= A/(A-A_0_)* is the relative area change of the GUV during tube elongation, *A_0_=4πR^2^* is the initial area of the GUV, *R* the GUV radius and *A=2πr_0_L*(*t*) is the area of the cylindrical tube; *r_0_* is the curvature radius of the tether and *L(t)* the tether length. Based on the considerations above, we described the dynamics of a sliding membrane with a one-point one-step master equation (Phillips et al., 2012; Van Kampen, 1992). The probability that the tip of a sliding membrane ends up at the site *n* on the MT lattice at the time *t+Δt* starting from the time *t* is the sum over all of the individual paths available to the system with the condition that the final state is fixed to one and can be written as:

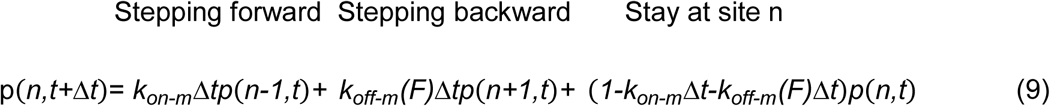

We transformed this equation into the more familiar continuum form as the function of the position *x* along the MT lattice, where the probability *p(x,t)* for finding the tip of the deformation at position *x* is equal to *p(n,t)* when *x=nd* as was done in (Phillips et al., 2012). This led to a result that represents a biased diffusion equation for the front of the sliding membrane of the deformation tip:

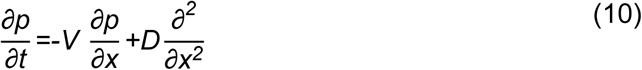

where

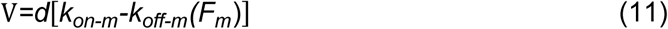

and

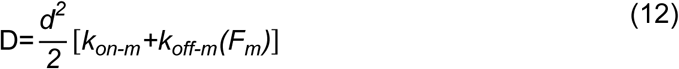

*V* is the average speed of the membrane tip and *D* is the variance of the tip position due to the stochastic nature of the process. As can be seen from Equations 7 and 11, V depends on *F*(*L*) or, in the context of tip’s probability function *p*, ultimately on *x*. To generate the dependence of speed for different time points of spreading (depicted in the Figures 5E right and S5B), we used iterative estimation of *x*(*t*) over time series with step *Δt*:

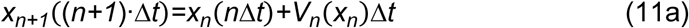

with the initial condition *x*(0)=0, which at the end provided values of *V_n_* for each time point *n Δt.* We chose the value of *Δt* = 0.5 s, and the variation in this parameter in the range of 0.1-5 s did not affect the final result. Curves in Figure 5B are averages of multiple solutions of Equation 11a with random values of GUVs physical parameters from Equations 7 and 8 described in the next section.

#### Parametrization of the model

We assumed that the affinity of the membrane for the MT is determined by the binding properties of the His-GFP-MTLS to MT. The value of the *k_off-m_ =* 1.7 s^-1^ was defined as the inverse of the dwell time of the His-GFP-MTLS on the MT lattice, obtained from the analysis of the His-GFP-MTLS single molecule binding events (Figures S5C-F). We generated the curves in Figure 5B with three different values of *k_on-m_* = (8,16,26) s^-1^. In our model, only *k_off-m_* was modified by the force required to pull a membrane tube as described by Equations 7 and 8.

The mechanical parameters of the membrane were obtained from the analysis of thermal fluctuations of the GUVs (Figure 3G,H). The precise value of the characteristic length of the potential barrier *ξ* between bound and unbound states is unknown for our system. However, the values reported in different systems are in the range between 1-2 nm (Nowak and Chou, 2010; Schnitzer et al., 2000). For modeling, we fixed *ξ* to 1 nm.

The radius of curvature of the deformation *r_0_* defines the number of MTLS molecules bound to the tip. In the 1D model *r_0_* was represented by a fixed length *L=dn_p_* in the range between 8 and 32 nm (Campàs et al., 2008)). For modeling, the number of MTLS molecules bound to the tip was randomly selected between 1 and 3.

The curves in Figure 5B are the average of 100 individual traces. For each trace, the lateral tension and the bending modulus of the simulated membrane were randomly sampled from a normal distribution with the mean and variance obtained from the analysis of the thermal fluctuations of the GUVs (Figure 3G,H). The radius *R* was sampled from a uniformly randomly distributed values ranging between 5 and 15 µm.

### Stochastic simulations

#### Membrane sliding along MT lattice

To simulate the dynamics of membrane tubes sliding along MTs, we have adapted the previously developed model for motors pulling fluid membranes (Campàs et al., 2008). His-GFP-MTLS molecules were represented by particles. MTs were represented by a discrete one-dimensional (single protofilament) lattices with *M* grid cells numbered from left to right (from the seed to plus end), and the membrane was described by a one-dimensional discrete lattice with *N* grid cells with *N<M,* also numbered from left to right (Figure 5D). The condition *N<M* guaranteed that the substrate for the adhesion was always available. Lattice spacing was fixed to the size of a tubulin dimer *d*=8 nm. Individual MT cells were considered attachment sites with capacity of 1 molecule and individual membrane sites were reservoirs of His-GFP-MTLS molecules. The number of His-GFP-MTLS molecules in a given cell at the MT grid represented the MT occupation number *n_MT_*. Similarly, the number of MTLS molecules occupying each site on the membrane was denoted as the membrane occupation number *n_m_*. Along the MT, His-GFP-MTLS molecule attachment sites were either empty (*n_MT_*=0) or occupied (*n_MT_*=1). We restricted the number of His-GFP-MTLS molecules attached to the MT at a given site to 1. However, along the membrane, each cell in the grid was unoccupied or occupied by several His-GFP-MTLS molecules.

The specified transitions of the His-GFP-MTLS molecules attached to a MT were:

1. Detachment from the MT with a rate *k_off-m_*. If the transition occurred at the tip of the deformation, the tube retracted to the next site containing a bound MTLS molecule.
2. Diffusion to the left or to the right with a rate *k_D-MT_*, if the site was empty.
3. Diffusion to the right from position *N* was not allowed.
4. Diffusion to the right from position 1 increased the number of proteins in the deformation by one.

The putative transitions of the His-GFP-MTLS molecules detached from a MT were:

5. Attachment to the MT with a rate *k_on-m_* if the site on the MT was unoccupied. If the transition happened at the tip, the deformation was extended by one site.
6. Diffusion to the left or to the right with a rate *k_D-M_* (Table 1, parameter 17) if the site was empty.

In the particular case of a membrane spreading along a MT lattice, the on-off rates along the deformation remained constant; however, at the tip of the deformation, the dissociation rate depended on the force barrier as explained above (see Equation 7).

#### Membrane spreading at the tips of growing MTs

The stochastic simulations describing the behavior of spreading membrane tubes in the presence of dynamic MTs included some additional features. As before, the MT was represented by a discrete 1D lattice, but with *N* grid cells numbered from left to right and the membrane was described by a discrete 1D lattice with *M* grid cells (Figure 6A). A feature of the simulation in the presence of growing MTs was the stochastic dynamics of the MTs. In the reaction scheme for MTs that was used in the simulations, MTs could be either in a growth phase or in a shrinkage phase.

For a MT in the growth phase, the possible events were:

1. Association of a tubulin dimer at position *N*: The length of the MT was increased by one dimer. The length of the tube was unchanged.
2. Dissociation of a tubulin dimer at position *N*: The length of the MT was decreased by one dimer. If the membrane was spreading up to the tip (M=N), it would also shrink to the next site containing a bound His-GFP-MTLS molecule.
3. Catastrophe: A change in the state from the growth phase to the shrinkage phase.
4. Rescue: A change in the state from the shrinkage phase to the growth phase.

The specified transitions of the His-GFP-MTLS molecules attached and detached from a MT were as before with the following special rule for His-GFP-MTLS molecules detached from a MT:

5. Attachment to the MT with a rate *k_on-m_* if the site on the MT was unoccupied. If the transition happened at the tip of the membrane, the deformation was extended by one site. If the membrane was spreading up to the tip (M=N) the transition was not allowed, because in our experiments, the membrane was never observed to spread beyond the MT tip.

As before, the dissociation rate was constant along the MT, but in the vicinity of the deformation tip, the rate depended on the force barrier. The association rate was regulated in space and time by the presence or lack of EB3.

### Boundary between the deformation and the vesicle

Position 1 in the grid characterized the connection of the deformation with the GUV (Campàs et al., 2008) and constituted the only source of His-GFP-MTLS molecules entering the membrane tether. We assumed that there is no influx of free His-GFP-MTLS molecules from solution. Since the linear density of His-GFP-MTLS molecules in the deformation must be constant over time, at this position the occupation numbers of the attached and detached molecules were kept constant and equal to 1 (*n_MT_*=*n_m_*=1). Diffusion from site 2 to 1 was not allowed, to keep the number of molecules in the deformation constant over time. The diffusion rate of the proteins that were not attached to the MT at position 1 was different from the other sites in the grid and was defined as *k_D-M1_*=*ρ_MTLS_*V_tube_*, where *ρ_MTLS_* is the linear density of the MTLS molecules in the deformation and *V_tube_* is the growth speed of the tube. This condition accounted for the influx of molecules entering the deformation due to the membrane flow resulting from the tube growth.

### Density of MTLS molecules in the stochastic simulation

At the beginning of the simulation, the linear density of the His-GFP-MTLS molecules was estimated as described previously(Campàs et al., 2008): *ρ_MTLS-linear_= ρ_MTLS-Dlinear_+ ρ_MTLS-Alinear_*, where *ρ_MTLS-Dlinear_* and *ρ_MTLS-Alinear_* were the linear density of the MTLS molecules detached and attached to the MT respectively. The linear density were defined as*: ρ_MTLS-Dlinear_=2π r_0_ ρ_MTLS-MT_* (*k_off-m_* /(*k_off-m_* + *k_on-m_*)) and *ρ_MTLS-Alinear_=2π r_0_ ρ_MTLS-MT_* (*k_on-m_* / (*k_off-m_* + *k_on-m_*)), *ρ_MTLS-MT_* was the surface density of MTLS molecules at GUVs surface in contact with MTs estimated experimentally, and *r_0_* was the curvature radius of the deformation.

### Simulation parameters in the absence of EB3

As before, the value for *k_off-m_* = 1.7 s^-1^ was defined as the inverse of the dwell time of His-GFP-MTLS molecules on MT lattice, obtained from the analysis of the single molecule binding events (Figure S5C-F). The stochastic rate of the detachment reaction was *aoff-m*= *k_off-m_*\**n_MT_*. As explained above, the dissociation rate remained constant along the MT, but was allowed to change at the very end of a membrane deformation. The first *n*_p_ MTLS molecules bound to the MT at the tip of a deformation shared the force to pull a tube. The parameter *n*_p_ was determined as a value of membrane occupancy *n_MT_* of the last four right elements. The parameters *σ_0,_κ* and *R* were random variables sampled from a normal distribution. The parameter *ξ* was fixed to 1 nm.

To determine the value of *k_on-m_* we compared the results obtained from the one-step model using different values for *k_on-m_* (Figures 5B and S5B). Based on the experimental data (Figure S5B), we found that *k_on-m_*=16 s^-1^ was a reasonable value to use in the stochastic simulations. This value of *k_on-m_* is the attachment rate of MTLS to MT lattice, which is constant along the MT and we assumed that was not affected by the tube force. In the absence of EB3, *k_on-m_* had the same value along the MT shaft and at the MT tip, because MTLS molecules at the front of the deformation cannot distinguish between MT shaft and MT tip. The stochastic rate of the attachment transition was *a_on-m_*= *k_on-m_*\**n_m_*.

### Simulation parameters in the presence of EB3

In the presence of EB3, the formation of a MTLS-EB3 complex imposed a spatial and temporal regulation on *k_on-m_*. The association rate of His-GFP-MTLS at MT tips that were not in contact with membranes was 2.5 fold higher than the on-rate of MTLS at the MT shaft (Figure S5F). We assumed that the kinetic constants of MTLS bound to the membrane displayed the same ratio, resulting in *k_on-m-tip_* = 2.5 * *k_on-m_*.

As mentioned above, every elemental stochastic reaction at a given position in the grid depends on the number of available molecules (occupation number). The number of EB3 molecules is high at the plus ends and decays towards the seed along the lattice of the MT, showing a comet-like shape. The distribution of the number of His-GFP-MTLS molecules measured from experiments showed the same pattern (Figures 1E and S1B). The stochastic rate of the attachment reaction to the tip of MTLS in the presence of EB3 can then be written as *a_on-m-tip_*= *k_on-m-tip_*\**n_m_*\**p_m-MTLS_*, where *p_m-MTLS_* is the occupation probability of MTLS along a MT starting from the MT tip measured experimentally (Figure 6C). MTLS molecules that detached from a MT diffused along the membrane with equal probability to the left or to the right. In the discretized representation, the diffusion of the proteins along the deformation is characterized by a constant rate *k_D-m_*=*D*_m_/*d^2^* (ref(Campàs et al., 2008)) with a stochastic rate *aD-m*= *k_D-m_*\**n_m_* where *D_m_* is the diffusion coefficient of the protein on the surface of the GUVs (Figure S5H,I).

Similarly, we considered symmetric diffusion of MTLS molecules bound to a MT, with the stochastic rate *a_D-MT_*= *k_D-MT_*\**n_MT_* and *k_D-MT_* =*D*_MT_/*d^2^*. Here, *D*_MT_ is the diffusion coefficient of MTLS molecules bound to the MT we considered that the diffusion rates were not affected by the presence of EB3 or by membrane forces.

### Gillespie algorithm: first-family method

The system was characterized by a rate matrix *k_S,F_* where *S=1,…N* indicates the sites in the 1D lattice representation and *F=1,…11* the number of potential transitions of the MTLS molecules. To implement the first-family method, we considered every transition *F* as a family with *S* potential reactions in each family(Gillespie, 2007). Every family *F* is then considered as a pseudoreaction with the stochastic rate 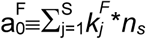, where 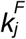 is the rate of the family *F* and *n_s_=*(*n_m_* or *n_MT_*) is the occupation number at the position *S*. To generate the time *t* to the next reaction event and the index pair (*F,S*) that identifies the type of transition and the position, we generated *F*+1 random numbers *r*_1_,…*r*_F+1._ We used the first *F* numbers to calculate

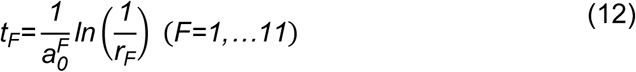

then

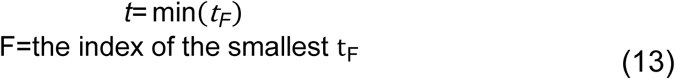

From the above, we determined the type of the next transition (detachment, attachment, diffusion, MT polymerization, MT depolymerization and MT transitions between growth and shrinkage). The position *S* of the transition in the grid was determined as:

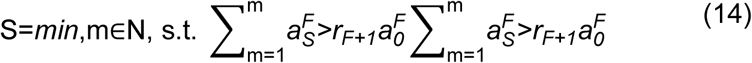

where 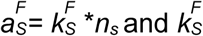 is the rate of the family *F* at position *S* on the grid. All the simulations were performed in MATLAB using custom routines

### Postprocessing of the simulation output

The output of each individual simulation was the position of the membrane tip (sliding along a MT shaft), or the position of the membrane and MT tip. We applied the following rules to sort the deformation events:

1. Events with a deformation of length lower than 200 nm were regarded as non-deformation events with zero length.
2. Evens classified as deformations (length>200 nm) were categorized as short tubes when the tube length was the range of 200-600 nm.
3. Tubes with a length >600 nm were considered as long deformations.
4. Only MTs that grew more than 200 nm were considered for analysis.
5. A detachment event happened when the distance between the membrane tip and the MT tip was longer than 200 nm.

## Data and software availability

All data that support the conclusions are available from the authors on request, and/or available in the manuscript itself. Data analysis was performed in MATLAB or using GraphPad Prism version 8.00 for (Mac OS X). The custom software used for data analysis and simulations in this manuscript can be found at https://github.com/RuddiRodriguez/; https://github.com/ekatrukha/KymoResliceWide. The software for the analysis of the GUVs thermal fluctuations was developed M. Mell, I. López Montero and R. Rodríguez-García at the Complutense University in the group of F. Monroy and is available at https://github.com/RuddiRodriguez/programnnn.

The computer code for simulations and analysis is available in https://github.com/RuddiRodriguez/Spreading;https://github.com/RuddiRodriguez/Spreading_membrane-MT_reaction_inside_EB.

Schematic representations were generated in Adobe Illustrator with the support of ChemDraw (PerkinElmer Informatics) and Biorender (©BioRender-biorender.com).

## Author Contributions

R.R.-G and V.A.V. designed and performed experiments and simulations, analyzed data and wrote the paper. M.P.L. and G.K. participated in developing the experimental setup, E.A.K. and L.C.K. contributed to data analysis and modeling, C.-Y. C., N.O., A.Ah., and M.O.S. contributed essential reagents. I.G. contributed to data collection and analysis, M.D and A.Ak. coordinated the project and wrote the paper.

## Acknowledgements

We thank M. Monti for developing a first version of a model for MT plus end-dependent force generation. This work was supported by the European Research Council Synergy grant 609822 to M.D. and A.Ak., the EMBO long-term fellowships to R.R.-G. and a grant from the Swiss National Science Foundation 31003A_166608 to M.O.S.

## Competing financial interests

The authors declare no competing financial interests.

## Legends to Supplementary Figures

**Figure S1, related to Figure 1.**
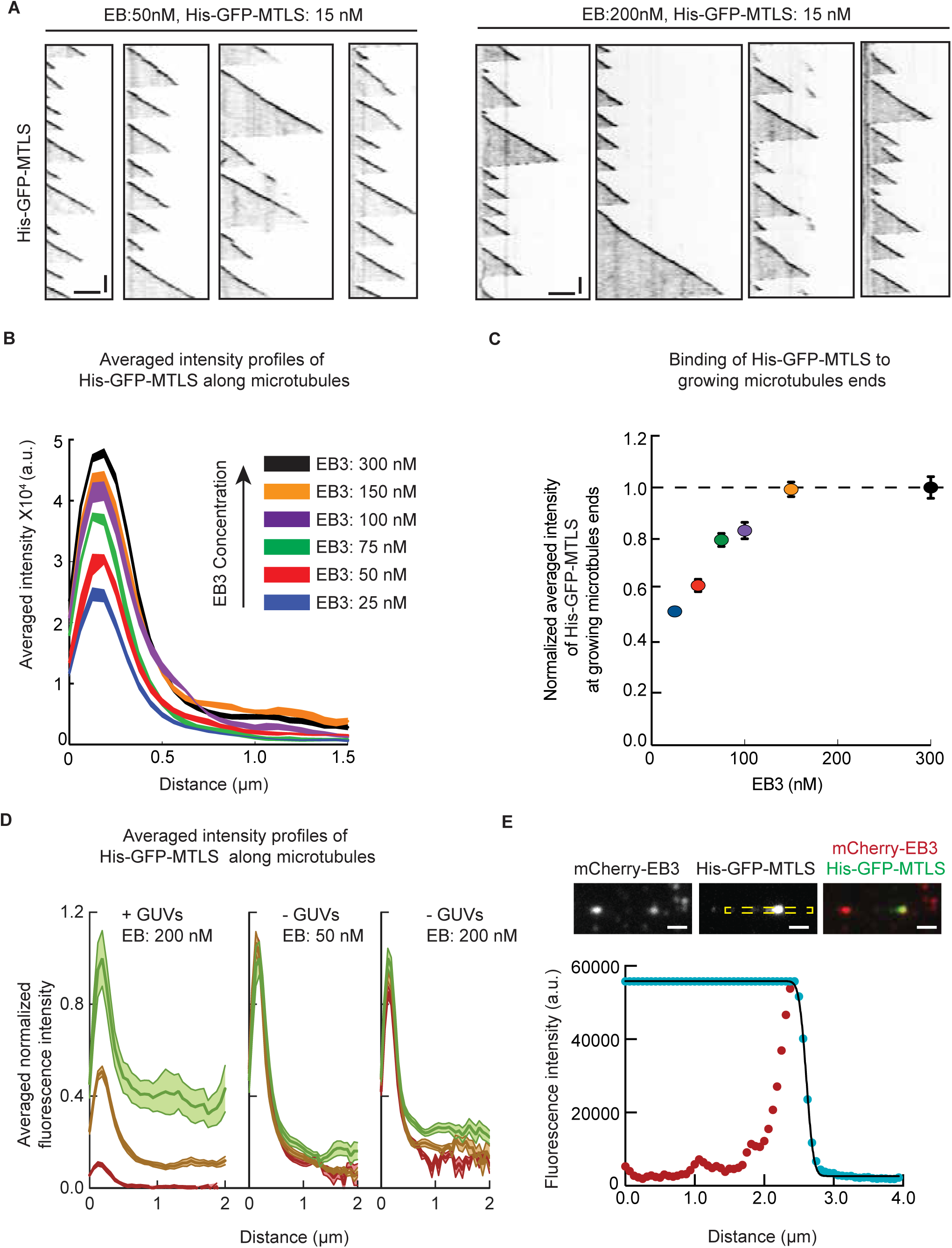
Analysis of the distribution of His-GFP-MTLS along MTs. (A) Representative kymographs illustrating the dynamics of MTs grown in presence of 15 nM of His-GFP-MTLS at different concentrations of EB3 (left, EB3 = 50 nM, right EB3=200 nM). Scale bars: horizontal (2 μm), vertical (60 s). (B) Averaged tip intensity profiles for His-GFP-MTLS in the presence of different EB3 concentrations and 20 μm of porcine tubulin. The shaded areas represent SEM. From low to high EB3 concentration, n=65, n=134, n=37, n=75, n=102, n=93. (C) Averaged maximum His-GFP-MTLS fluorescence intensities at MT plus ends as a function of EB3 concentration. Data were normalized to the average maximum intensity obtained at the highest EB3 concentration. The color codes and n numbers are the same as in panel (B). (D) Normalized averaged tip intensity profiles for His-GFP-MTLS along MTs at different times in presence or in absence of GUVs. The profiles were normalized to the maximum intensity at time corresponding to the first movie acquired after the addition of the GUVs (t=0). In presence of GUVs t=0, n=17; t=10min, n=25; t=20min, n=18. In absence of GUVs EB3:50, t=0, n=28, t=30 min, n=39, t=45 min, n=23. EB3:50, t=0, n=15, t=15 min, n=51, t=30 min, n=33. (E) Top: Snapshot of a TIRFM time lapse movie. Scale bar: 1 μm. Bottom: MT tip intensity profile (red dots) and the same profile after the maximum intensity was assigned to each of the preceding points along the MT (blue dots). The transformed profile was fitted with the error function (black line). Figure illustrated position fitting for consecutive the intensity profiles averaging in Figure S1A,C.

**Figure S2, related to Figure 2.**
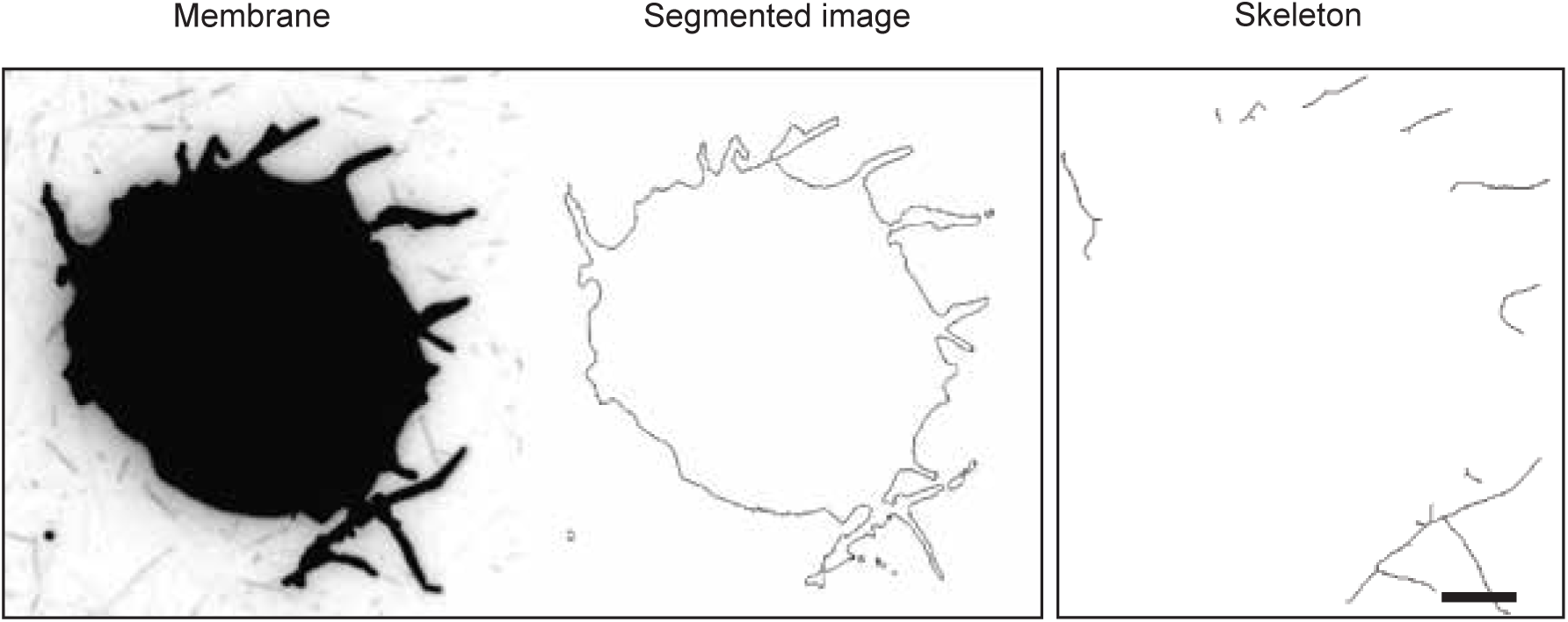
Analysis of membrane tubulation. Left: Snapshot of a tubular membrane network. The image is the maximum intensity projection obtained from 100 frames of time lapse images. Center: Segmented tubular network. Right: Skeletal representation of the tubular network. Scale bar: 5 μm.

**Figure S3, related to Figure 3.**
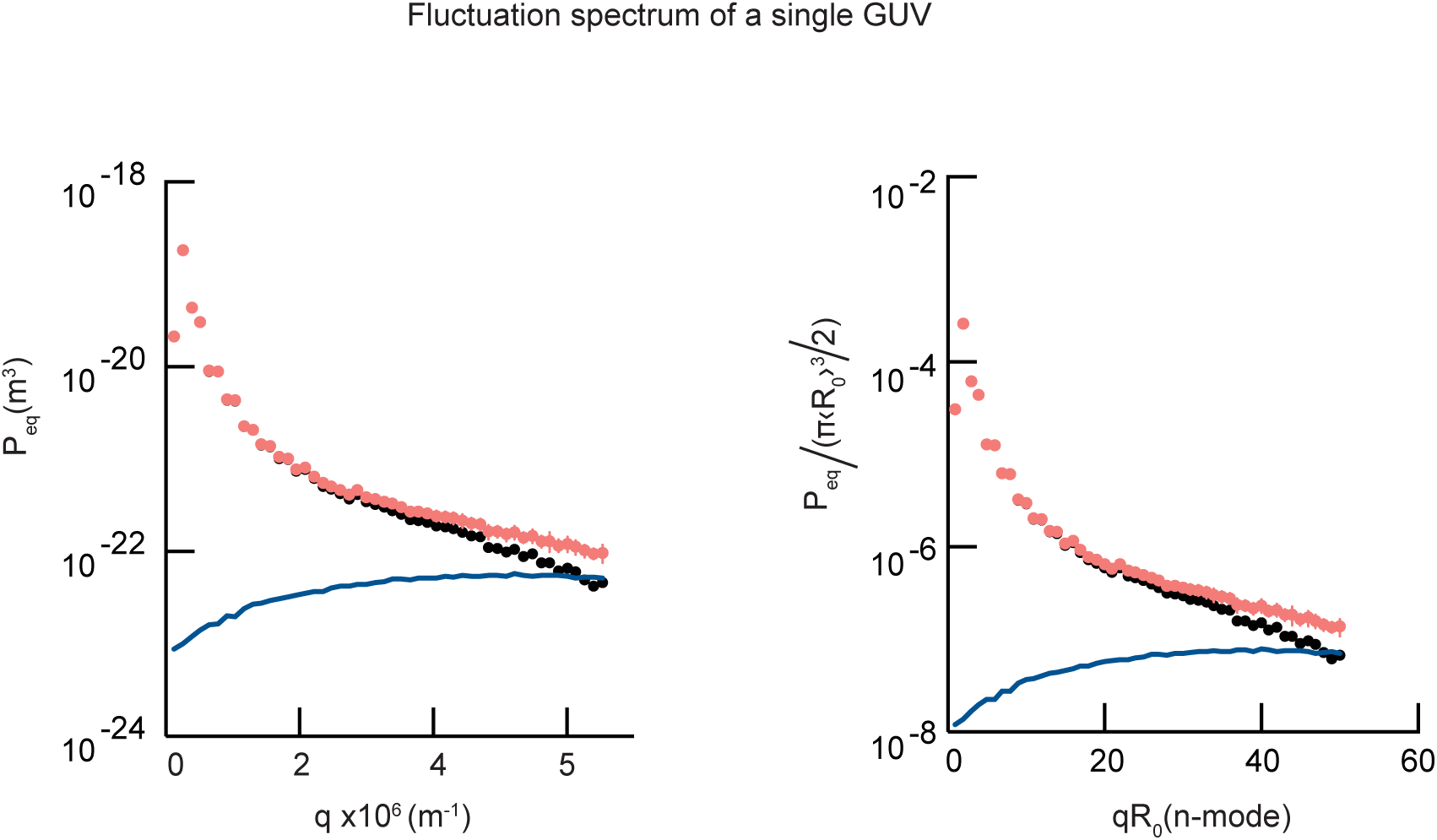
Fluctuation spectrum of a GUV. (Left) Experimental fluctuation spectrum calculated from the time-averages of the quadratic fluctuation amplitudes of the equatorial modes (red dots) measured for a single GUV with a radius of 10 µm. The blue line is the systematic contribution of the pixelization noise to the experimental fluctuation amplitude, which increases with the mode *n*. Black dots represent the amplitudes of fluctuations corrected for the pixelization noise. (Right) The spectrum was normalized by the factor *A=*π〈*R_0_*〉*^3^*/*2*. Error bars represent SE.

**Figure S4, related to Figure 4.**
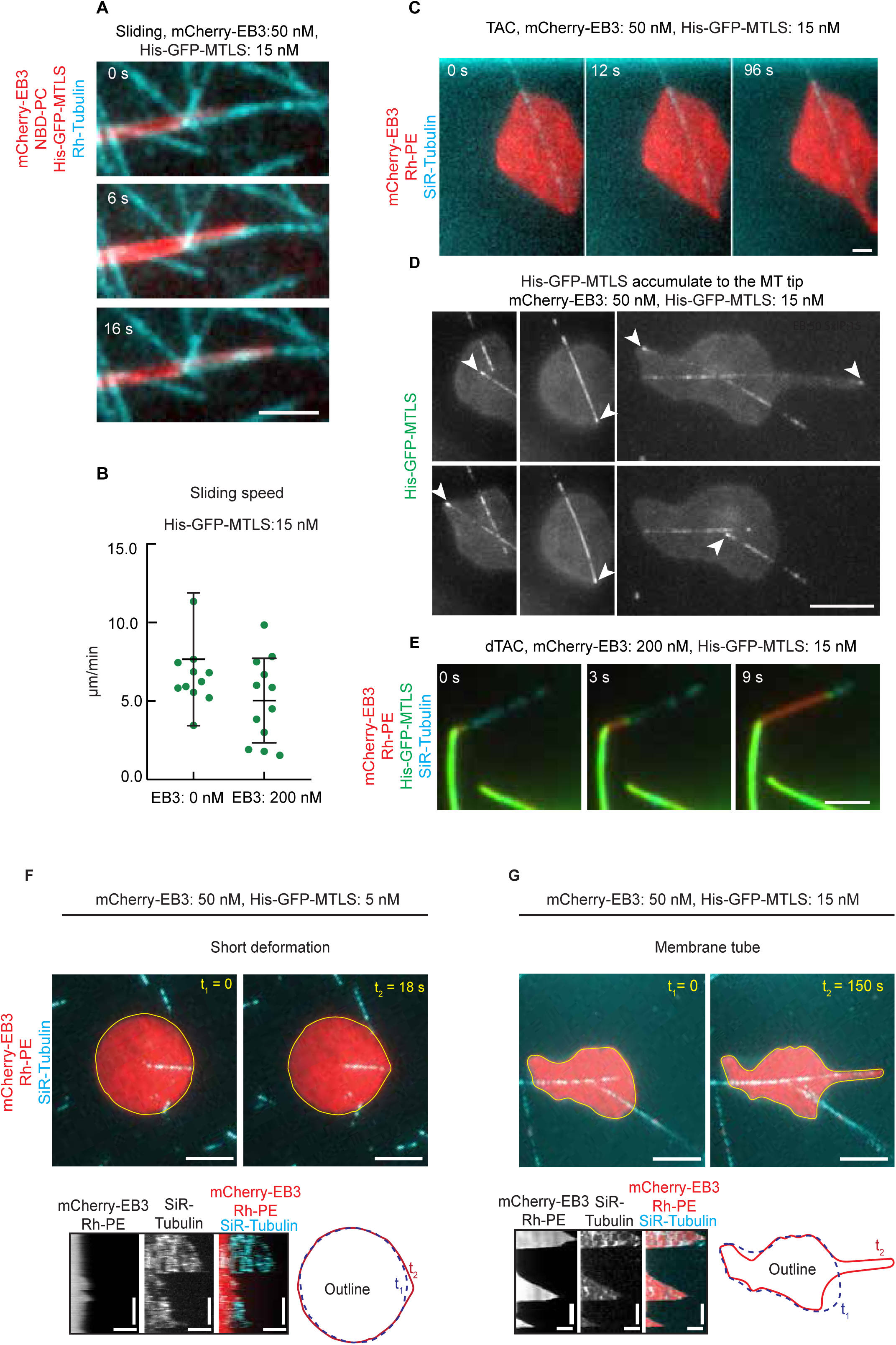
Three mechanisms of MT-induced membrane tube formation. (A,C,E) Time lapse images of a membrane tube moving along a MT shaft (sliding, A); together with the plus end of a growing MT (TAC, B) or by attachment to the plus end of a depolymerizing MT (dTAC, E). Scale bar: 2 μm. (B) Speed of membrane sliding at 0 nM (n=9), and 200 nM (n=12) EB3 and 15 nM His-GFP-MTLS. Error bars indicate SD. (D) Snapshot of a GUVs in contact with the MTs. The white arrow head shows the accumulation of the His-GFP-MTLS protein at the tip of the MTs. Only the His-GFP-MTLS channel is showed. Scale bar: 5 μm. (F,G) Snapshots (top) and the corresponding kymographs (bottom) of a MT deforming a GUV at 50 nM mCherry-EB3 and 15 nM His-GFP-MTLS. GUV contours are shown by yellow lines. (F) The membrane detaches from the MT before developing a tubular shape. (G) The membrane remains attached to the MT tip during growth and shrinking phases. Scale bars: 5 μm (top panels), horizontal, 0.5 μm, vertical, 2 min (kymograph in (F)); horizontal, 2 μm, vertical, 2 min (kymograph in (G)).

**Figure S5, related to Figure 5.**
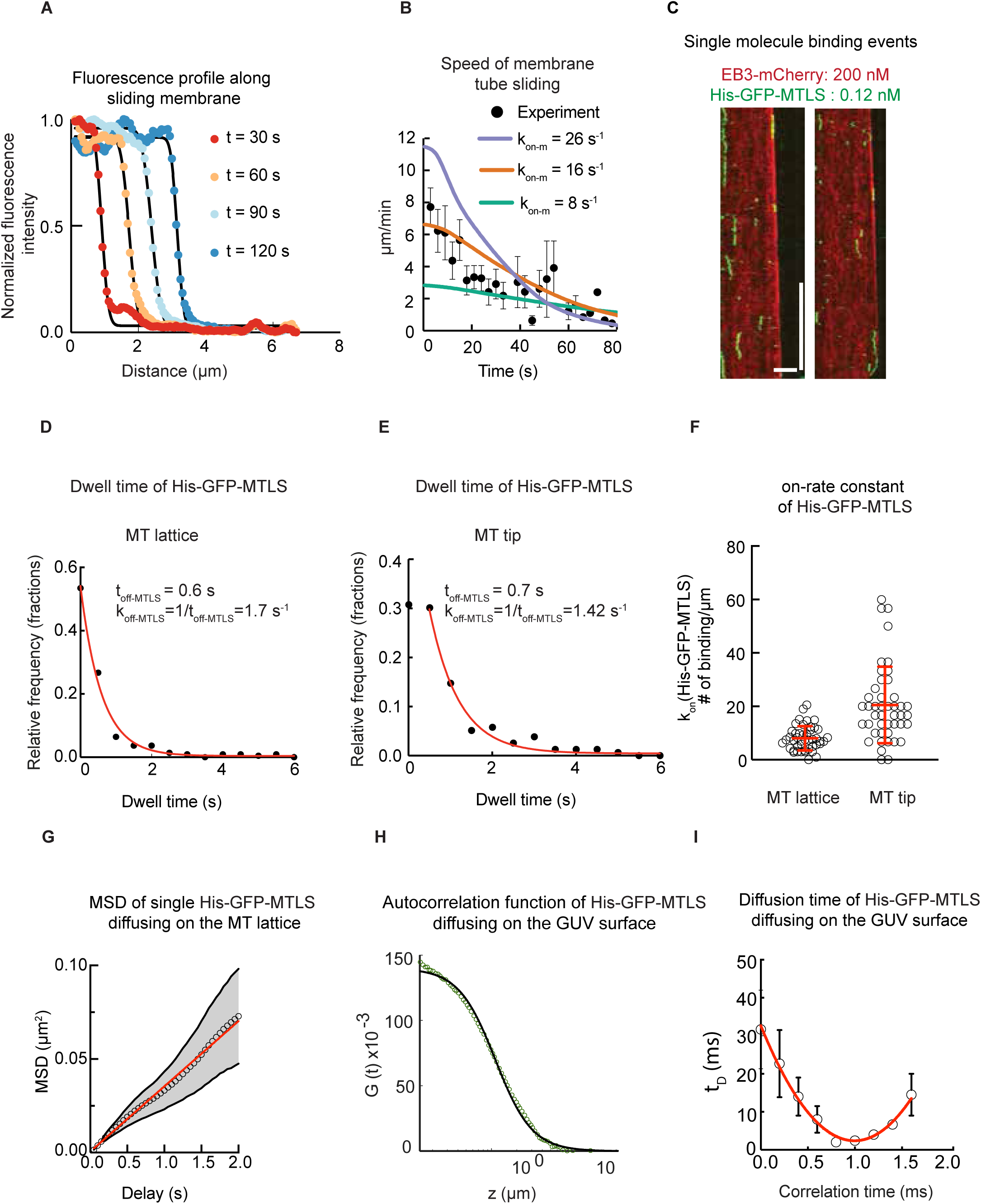
Determination of parameters used for modelling and stochastic simulations. (A) Membrane tip intensity profile at different time points measured from a GUV sliding along a MT shaft in presence of 15 nM of His-GFP-MTLS without EB3. The position of the membrane moving along a MT shaft was determined from the fitting of the intensity profiles to the step function. (B) Comparison between curves representing the speed of membrane tube sliding along MT lattice modelled using different association rates and the experimentally determined membrane sliding speeds. (C) Kymographs of dynamic MTs grown in presence of 0.2 nM His-GFP-MTLS and 200 nM of mCherry-EB3. (D-E) Histograms of dwell time of single His-GFP-MTLS molecules at MT lattice (C) and growing MT plus ends (D); experimental conditions were the same as in panel (C). The red lines represent monoexponentially fits. (F) Association rate of the His-GFP-MTLS on MT lattice and growing MT plus ends, expressed in number of binding events per length. (G) Mean square displacement (MSD) for single His-GFP-MTLS molecules diffusing on MT lattice. (H) Representative autocorrelation function obtained from the diffusion of the membrane-bound His-GFP-MTLS molecules. (I) Diffusion time of the membrane-bound His-GFP-MTLS molecules measured by FCS as a function of the position in z.

## Legends to Videos

**Video S1. In vitro Plus-End Tracking of His-GFP-MTLS in presence of EB3.** The sample was prepared with 15 nM of His-GFP-MTLS, 50 nM of EB3 (left) and 200 nM of EB3 (right). Images were acquired sequentially using TIRFM at with a 3 s interval. The movie is displayed at 5 fps. Scale bar: 5 μm.

**Video S2**. **Membrane tubes growing from a GUV in the presence of MTs.** Images of Rh-PE (red, membrane), GFP (green, His-GFP-MTLS), mCherry-EB3 (red) and SiR (cyan, MT) tubulin were acquired sequentially using TIRFM at with a 3 s interval. The movie is displayed at 5 fps. Scale bar: 5 μm.

**Video S3**. **Membrane tube sliding along a MT.** Images of Rh-PE (red, membrane), GFP (green, His-GFP-MTLS), mCherry-EB3(red) (left) and SiR tubulin (MT) (right) were acquired sequentially using TIRFM at with a 3 s interval. The movie was displayed at 5 fps. Scale bar is 2 μm.

**Video S4**. **Membrane tube extending together with a growing MT plus end.** Images of Rh-PE (red, membrane), GFP (green, His-GFP-MTLS), mCherry-EB3 (red) (left) and SiR tubulin (MT) (right) were acquired sequentially using TIRFM at with a 3 s interval. The movie is displayed at 5 fps. Scale bar: 2 μm.

**Video S5**. **Membrane tube pulled by a depolymerizing MT.** Images of Rh-PE (red, membrane) and GFP (green, His-GFP-MTLS) (left) and SiR tubulin (MT) (right) were acquired sequentially using TIRFM at with a 3 s interval. Note the strong accumulation of the His-GFP-MTLS at the tip of the membrane deformation. The movie is displayed at 1 fps. Scale bar: 2 μm.

**Video S6. Extension and retraction of MT-dependent non-tubular GUV deformations.** Images of Rh-PE (red, membrane), mCherry-EB3 (red) and SiR tubulin (cyan, MT) were acquired sequentially using TIRFM at with a 3 s interval. The movie is displayed at 5 fps. Scale bar: 3 μm.

**Video S7**. **Qdot transport with the tip of the growing MT**. Three channels were imaged by TIRFM sequentially at 1.3 s interval: Qdot705-streptavidin (red), HiLyte488-tubulin (cyan), Bio-mCherry-MTLS (15 nM, green) and mCherry-EB3 (100 nM, green). The corresponding kymograph is shown in Fig 7B. The movie is displayed at 30 fps. Scale bar: 5 μm.

**Video S8**. **A growing MT end pushes a trapped glass bead**. A 1 μm glass bead coated with bio-mCherry-MTLS trapped in a soft trap was imaged using DIC microscopy at 8 fps and every 10 frames were averaged. The movie is displayed at 30 fps. The QPD trace corresponding to this experiment is shown in Fig 7e.

## Notes

#### Summary of Updates

This version of the manuscript was revised to improve the clarity of the presentation.

